# Development of force-field corrections for the RNA A-bulge motif

**DOI:** 10.64898/2026.08.26.747445

**Authors:** Takafumi Kudo, Toru Ekimoto, Tsutomu Yamane, Mitsunori Ikeguchi

**Author notes:** Corresponding Author: Mitsunori Ikeguchi.

## Abstract

Many functional RNA motifs adopt structures that deviate from the canonical A-form helix and are emerging targets for RNA-directed therapeutics. The microtubule-associated protein tau (MAPT) A-bulge motif (5**′**-GCAGU/5**′**-ACGU) is one such motif. Because its structure is stabilized by a delicate balance of local interactions, its accurate modeling remains a major challenge for molecular dynamics (MD) simulations. The experimentally determined nuclear magnetic resonance (NMR) structure of the MAPT A-bulge motif provides a stringent test of whether RNA force fields can accurately reproduce the experimentally observed conformation. Most current AMBER-family RNA force-field models have incorrectly favored a non-native base-triple state of the MAPT A-bulge motif over the experimentally observed stacked state. Structural comparison of the stacked and base-triple conformations revealed that overly favorable NH**□**–N hydrogen bonds between the bulged adenosine and an adjacent Watson–Crick base pair were the primary source of this imbalance. We developed gHBfix-18Ab, an 18-component hydrogen-bond correction that distinguishes NH and NH**□** donors. gHBfix-18Ab was combined with the previously developed OL3_CP_ and NBfix_0BPh_ corrections to generate the composite model gHBfix-18Ab*. This model restored the experimentally observed stacked state as the global minimum in the calculated free-energy profile and improved agreement with NMR-derived distance data for the A-bulge region. Importantly, gHBfix-18Ab* did not produce marked structural destabilization of the cUUCGg tetraloop, a widely used benchmark for RNA force-field validation, suggesting that the refinement preserves the stability of the unrelated RNA motif. These results demonstrate that targeted refinement of hydrogen-bond interactions provides a practical strategy for systematic improvement of RNA force fields toward more accurate modeling of noncanonical RNA motifs.

**Graphical Summary:** 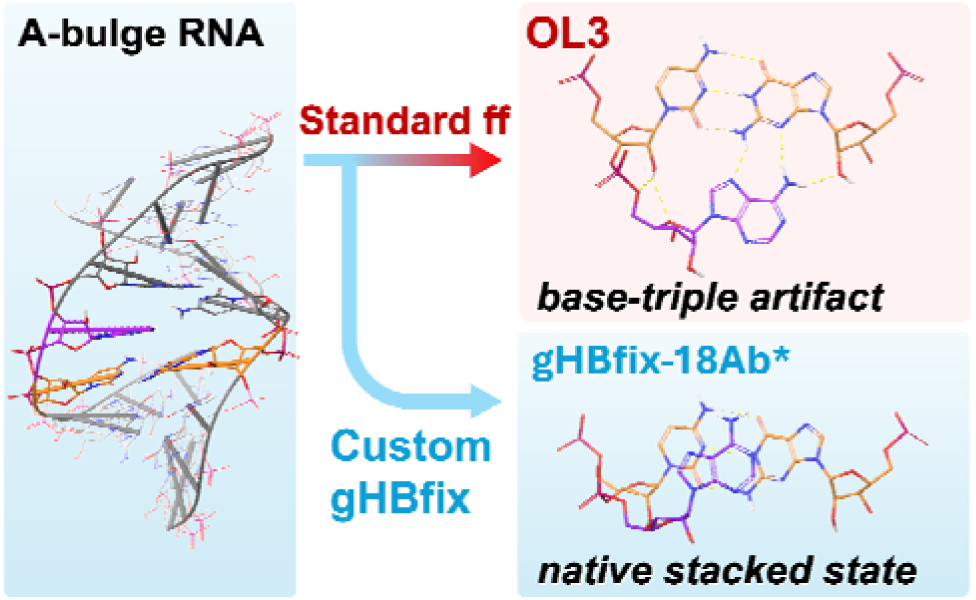

## 1. Introduction

RNA molecules regulate diverse biological processes through sequence-dependent structural motifs, many of which are emerging targets for RNA-directed therapeutics. Sequence-dependent local interactions stabilize distinct conformations and define the equilibrium among competing conformational states despite the intrinsic flexibility of RNA molecules. In particular, noncanonical RNA motifs adopt structures that deviate from the canonical A-form helix and are often stabilized by a delicate balance of local interactions, making accurate modeling of their conformational ensemble a major challenge for molecular dynamics (MD) simulations. The reliability of RNA MD simulations therefore depends critically on the force-field model used to describe these conformational equilibria. Thus, evaluating RNA force fields is important not only for their ability to retain experimentally observed conformations but also for their ability to reproduce the relative stability of experimentally observed and competing non-native states.^1^ Adenosine bulges (A-bulges) are compact noncanonical RNA motifs containing an unpaired adenosine within an otherwise Watson–Crick base-paired helical region. Because experimentally determined structures are available for several A-bulges, these motifs provide useful benchmarks for assessing how accurately RNA force fields reproduce the balance between native and competing conformational states.

One such example is the A-bulge motif from the MAPT pre-mRNA exon 10 splicing regulatory element (5**′**-GCAGU/5**′**-ACGU). Its nuclear magnetic resonance (NMR) structure adopts a stacked conformation in which the bulged A6 residue lies between the adjacent C5 and G7 bases (Figure 1).^2^ However, an AMBER RNA force field with revised *χ* and *α/γ* torsional parameters instead overstabilizes a base-triple state, in which A6 forms hydrogen bonds with the adjacent C5–G17 Watson–Crick base pair (Figure 2B).^3^ Because the stacked and base-triple states differ primarily in the position and hydrogen-bonding pattern of the bulged adenosine, the MAPT A-bulge provides a useful test case for evaluating how RNA force fields balance an experimentally supported stacked state against a competing base-triple state.

**Figure 1.**
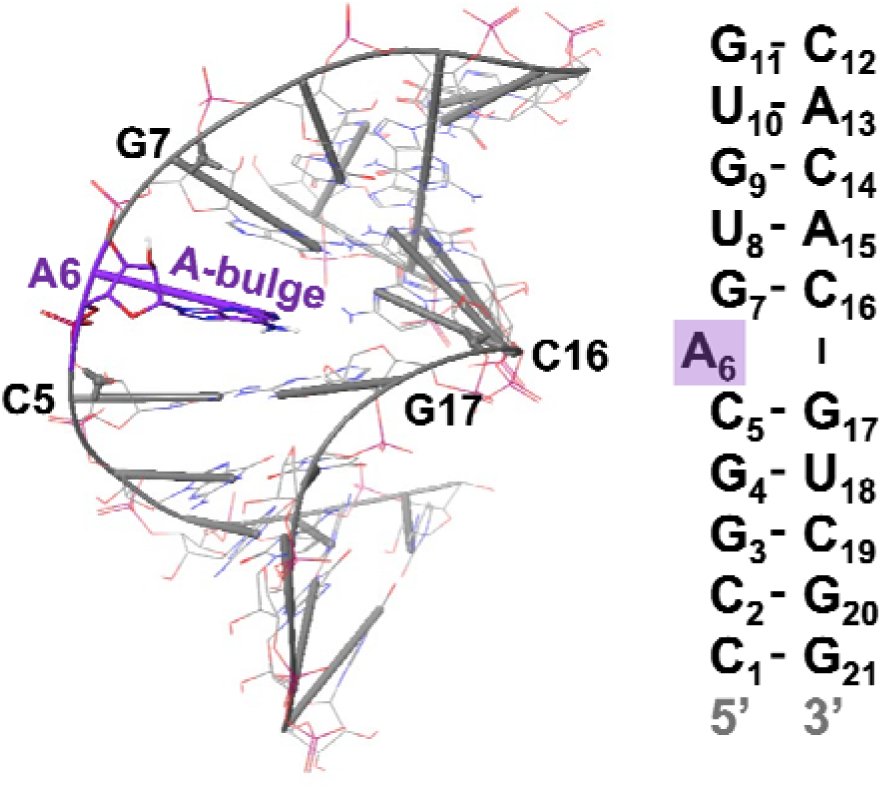
Sequence and NMR structure of the MAPT pre-mRNA A-bulge motif used in this study. The sequence corresponds to the RNA construct used for the NMR structure (Protein Data Bank (PDB) ID: 6VA1).^2^ The bulged adenosine residue A6 is highlighted, and the residue defining the A-bulge motif are indicated in the structural model.

**Figure 2.**
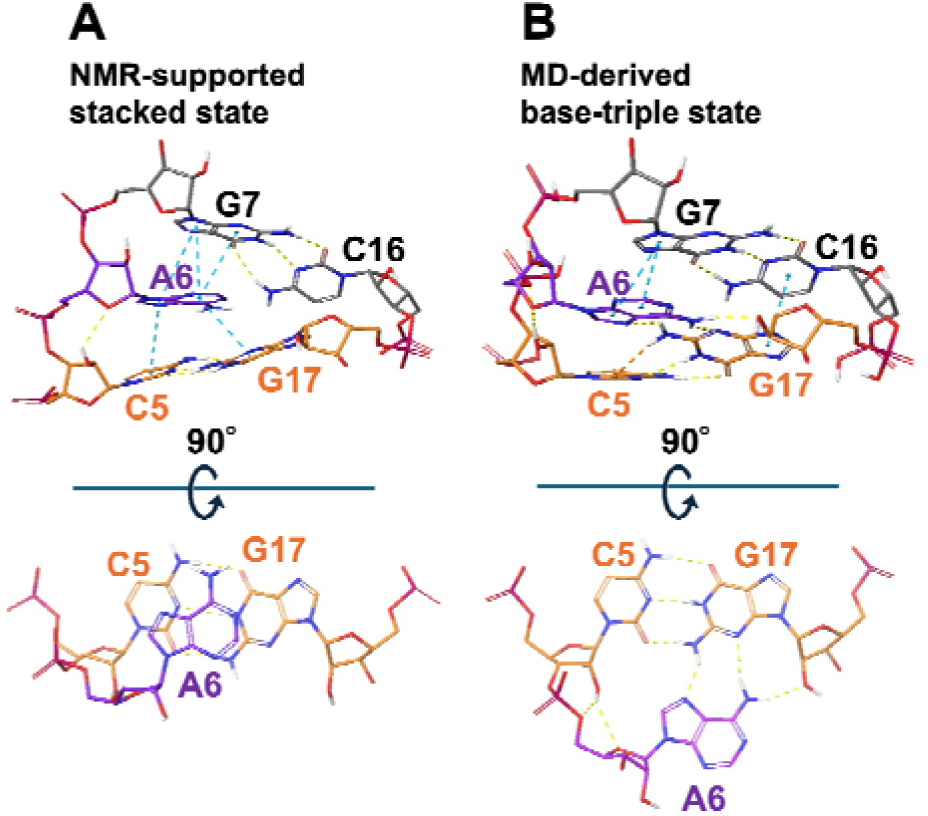
Structural comparison of the stacked and base-triple conformations of the MAPT A-bulge motif. (A) Experimentally supported stacked conformation based on the NMR structure (PDB ID: 6VA1).^2^ (B) Base-triple conformation extracted from our enhanced-sampling simulation with OL3 force field. Yellow dashed lines indicate hydrogen bonds, and blue dashed lines indicate π–π stacking interactions.

Among current RNA force fields, AMBER-family force fields have been widely used, and successive refinements have been introduced to address known limitations in their description of RNA conformations and interactions. Building on the widely used OL3 force field, ^4–7^ refinements have targeted several classes of force-field terms, including torsional parameters, phosphate and other nonbonded interactions, backbone corrections, and hydrogen bonding (Table 1). These include OL3_CP_, which incorporates modified phosphate-oxygen Lennard–Jones parameters developed to improve phosphate solvation;^8^ NBfix_0BPh_, which adjusts pair-specific Lennard–Jones parameters for intranucleotide H8···O5**′** and H6···O5**′** contacts to relax steric clashes;^9^ and gHBfix21, which applies donor/acceptor-type-specific corrections to selected intramolecular hydrogen bonds.^10,11^ However, it remains unclear whether these refinements correctly reproduce the balance between the experimentally observed stacked state and the competing non-native base-triple state of the MAPT A-bulge.

**Table 1.**
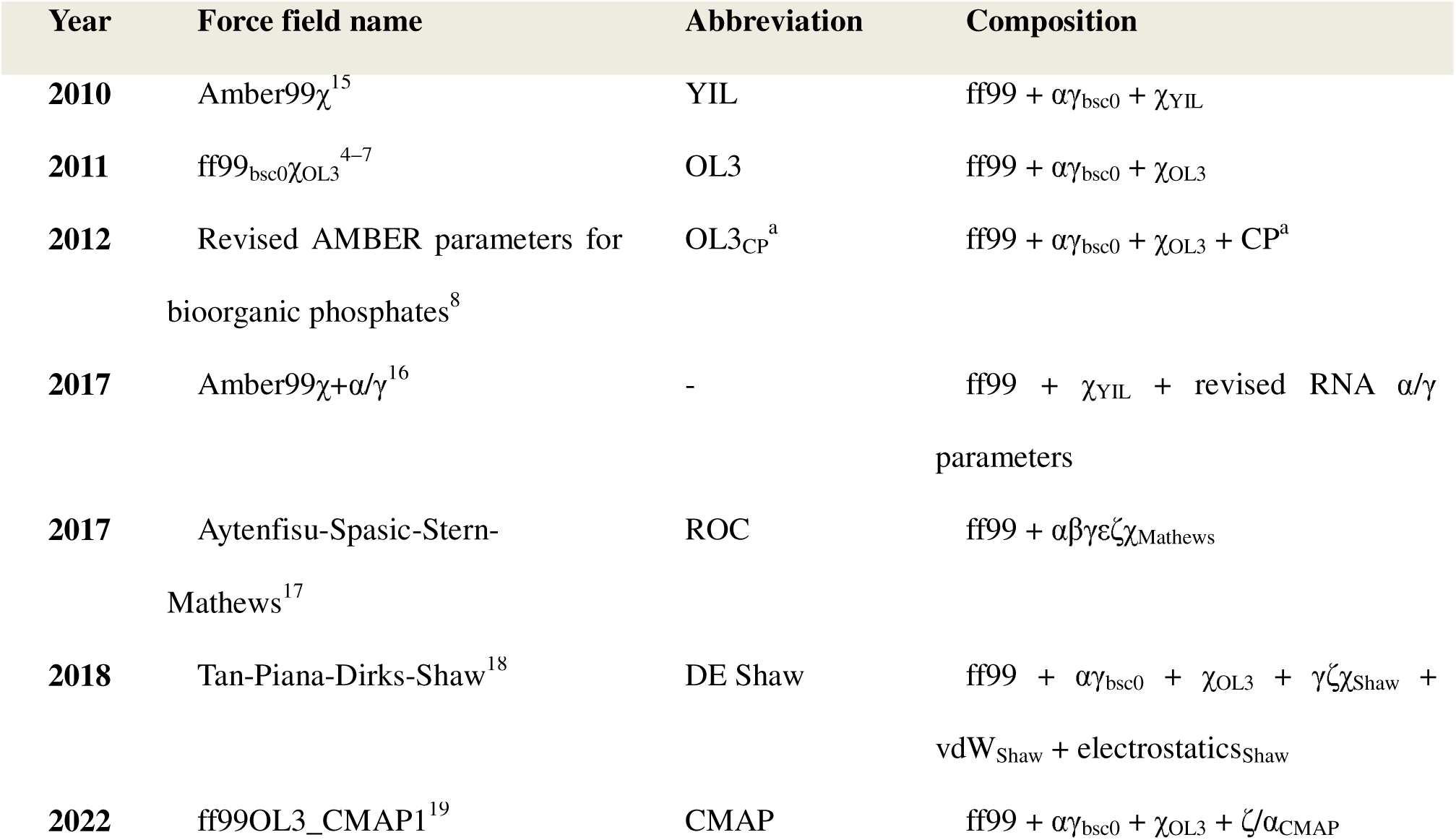
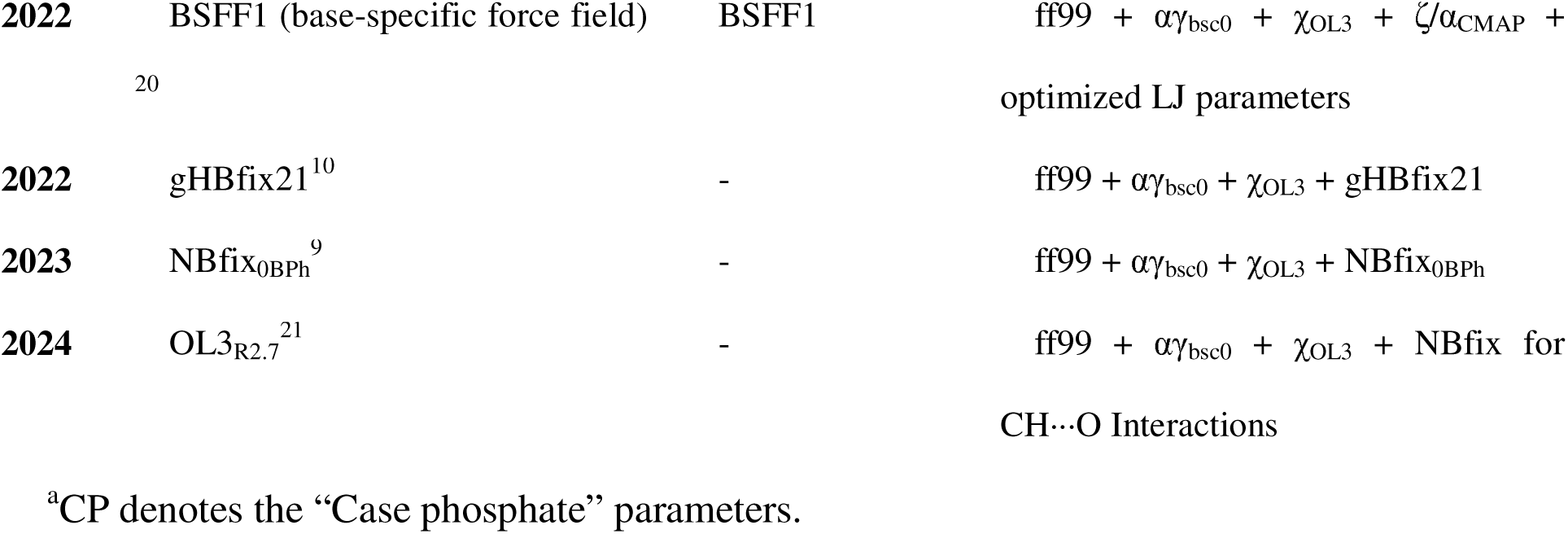
Representative AMBER-family RNA force fields and their refinements. ^4–10,15–21^

In this study, we aimed to identify RNA force-field conditions that reproduce the experimentally observed stacked state of the MAPT A-bulge while avoiding excessive stabilization of the competing base-triple state. We first evaluated representative AMBER-family RNA force fields by comparing the stacked/base-triple conformational equilibrium using enhanced-sampling free-energy calculations together with agreement with NMR-derived nuclear Overhauser effect (NOE) distance restraints.^3,12,13^ We compared local interactions in the stacked and base-triple conformations to identify those responsible for the conformational imbalance. This guided the development of gHBfix-18Ab, an 18-component hydrogen-bond correction that distinguishes NH and NH**□** donor groups, allowing their interactions to be tuned independently. The suffix “Ab” denotes A-bulge. We further evaluated gHBfix-18Ab in combination with the previously developed OL3_CP_ and NBfix_0BPh_ corrections,^8,9^ resulting in the composite force-field model gHBfix-18Ab*. Finally, we evaluated gHBfix-18Ab* using the cUUCGg tetraloop (Protein Data Bank (PDB) ID: 2KOC), a well-established RNA benchmark, to assess whether the refinement preserved the structural stability of this unrelated RNA motif.^14^

## 2. Methods

### 2.1. RNA Simulation Systems

The initial structure of the MAPT pre-mRNA A-bulge system was prepared from the first model in the NMR structure of the MAPT pre-mRNA exon 10 splicing regulatory element (5′-GCAGU/5′-ACGU; PDB ID: 6VA1).^2^ All nucleobases retained their standard protonation states as confirmed using the Epik module at pH 7.4 in Schrödinger Maestro version 2022-3.^22,23^ The simulation systems were constructed and solvated using the Solution Builder module of CHARMM-GUI.^24–26^

For solvation, the Optimal Point Charge (OPC) water model^27^ was used for the OL3, YIL, ROC, CMAP, OL3_CP_, gHBfix21, and gHBfix-18Ab-based systems, whereas the TIP4P-D water model^28^ was used for the DE Shaw force field. An octahedral simulation box was used for all systems. Na**□** and Cl**□** ions were added to neutralize the total system charge and to reach a final NaCl concentration of 0.15 M.

The same system-construction procedure was applied to the cUUCGg tetraloop system, which was prepared from the NMR structure (PDB ID: 2KOC).^14^ The force-field models, correction schemes, simulation systems, and simulation protocols used for the two systems are summarized in Table 2.

**Table 2.**
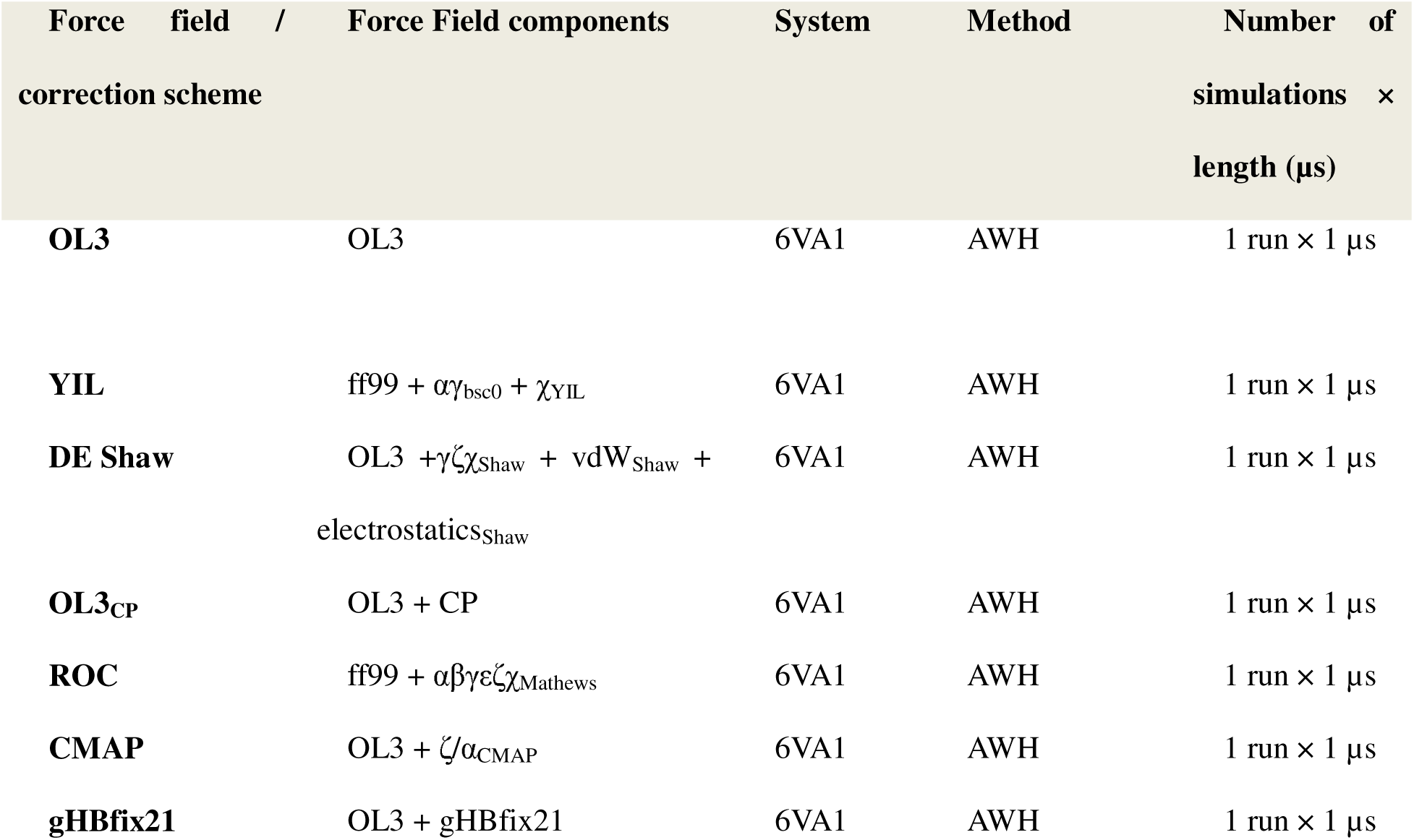
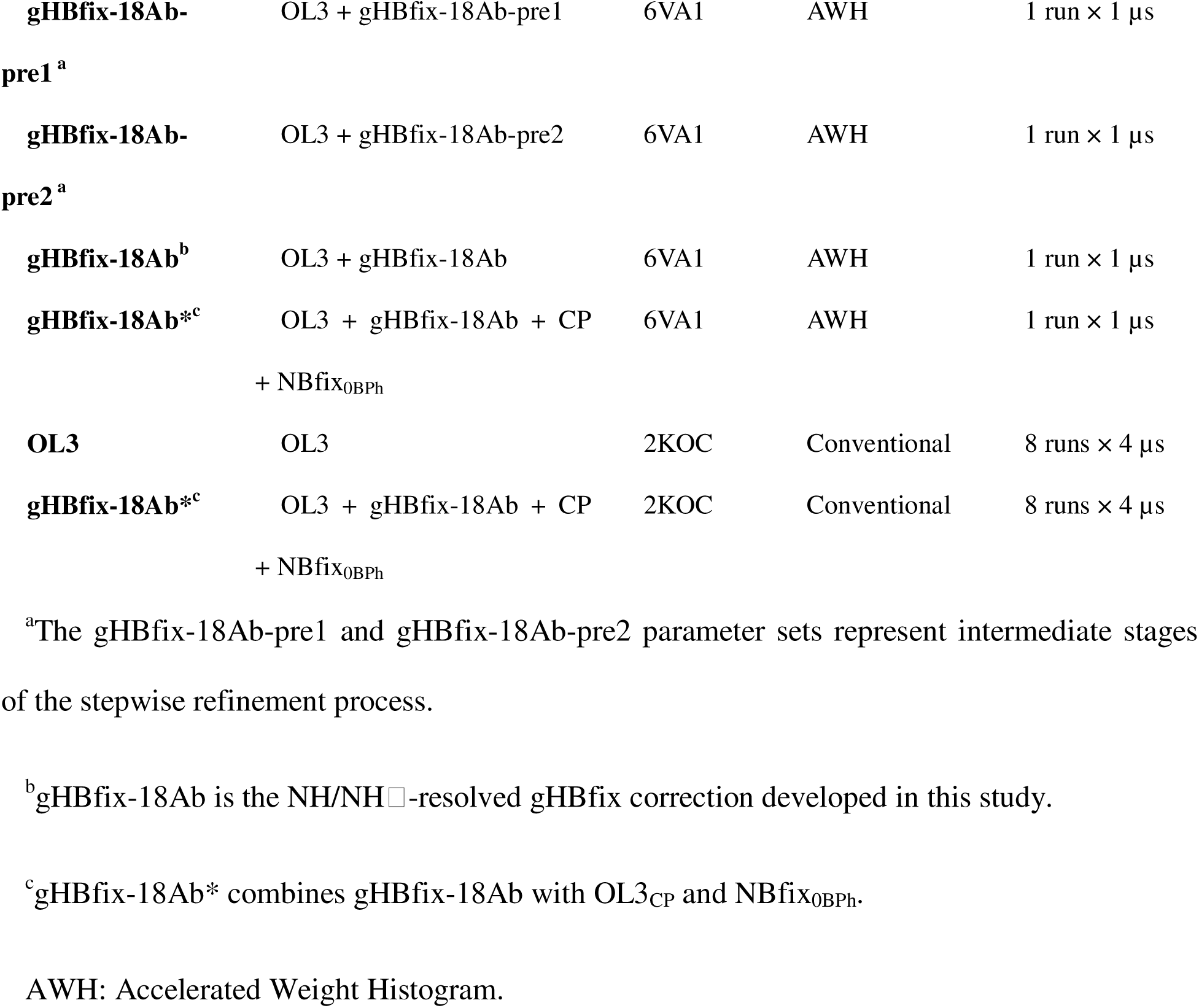
Summary of MD simulations performed in this study.

| Force field /<br>correction scheme | Force Field components | System | Method | Number of<br>simulations ×<br>length (μs) |
| --- | --- | --- | --- | --- |
| **OL3** | OL3 | 6VA1 | AWH | 1 run × 1 μs |
| **YIL** | ff99 + $\alpha\gamma_{\text{bsc0}}$ + $\chi_{\text{YIL}}$ | 6VA1 | AWH | 1 run × 1 μs |
| **DE Shaw** | OL3 + $\gamma\zeta\chi_{\text{Shaw}}$ + $\text{vdW}_{\text{Shaw}}$ +<br>electrostatics<sub>Shaw</sub> | 6VA1 | AWH | 1 run × 1 μs |
| **OL3<sub>CP</sub>** | OL3 + CP | 6VA1 | AWH | 1 run × 1 μs |
| **ROC** | ff99 + $\alpha\beta\gamma\epsilon\zeta\chi_{\text{Mathews}}$ | 6VA1 | AWH | 1 run × 1 μs |
| **CMAP** | OL3 + $\zeta/\alpha_{\text{CMAP}}$ | 6VA1 | AWH | 1 run × 1 μs |
| **gHBfix21** | OL3 + gHBfix21 | 6VA1 | AWH | 1 run × 1 μs |
| **gHBfix-18Ab-pre1<sup>a</sup>** | OL3 + gHBfix-18Ab-pre1 | 6VA1 | AWH | 1 run × 1 μs |
| **gHBfix-18Ab-pre2<sup>a</sup>** | OL3 + gHBfix-18Ab-pre2 | 6VA1 | AWH | 1 run × 1 μs |
| **gHBfix-18Ab<sup>b</sup>** | OL3 + gHBfix-18Ab | 6VA1 | AWH | 1 run × 1 μs |
| **gHBfix-18Ab\*<sup>c</sup>** | OL3 + gHBfix-18Ab + CP<br>+ NBfix<sub>0BPh</sub> | 6VA1 | AWH | 1 run × 1 μs |
| **OL3** | OL3 | 2KOC | Conventional | 8 runs × 4 μs |
| **gHBfix-18Ab\*<sup>c</sup>** | OL3 + gHBfix-18Ab + CP<br>+ NBfix<sub>0BPh</sub> | 2KOC | Conventional | 8 runs × 4 μs |

<sup>a</sup>The gHBfix-18Ab-pre1 and gHBfix-18Ab-pre2 parameter sets represent intermediate stages of the stepwise refinement process.
<sup>b</sup>gHBfix-18Ab is the NH/NH □ -resolved gHBfix correction developed in this study.
<sup>c</sup>gHBfix-18Ab\* combines gHBfix-18Ab with OL3<sub>CP</sub> and NBfix<sub>0BPh</sub>.
AWH: Accelerated Weight Histogram.

### 2.2. Construction of Force-Field Models

The OL3,^5–7^ YIL,^15^ and DE Shaw^18^ systems were generated using the corresponding force-field options in CHARMM-GUI Solution Builder. For the OL3_CP_,^8^ ROC,^17^ and CMAP^19^ models, the CHARMM-GUI-generated OL3 coordinates were retained as a common structural template, and the topology and force-field parameters were regenerated with AmberTools23^29^ tLEaP. For OL3_CP_ and ROC, the corresponding AmberTools leaprc command files (leaprc.RNA.LJbb and leaprc.RNA.ROC, respectively) were used during topology generation. In the CMAP model, the ff99OL3_CMAP1 parameter files obtained from the developers’ repository were loaded in tLEaP to generate the corresponding AMBER topology. The resulting AMBER topology files and the coordinate files were converted to GROMACS input files using ParmEd.^29^

NBfix_0BPh_ is a pair-specific Lennard–Jones correction for intranucleotide base–phosphate contacts, corresponding to purine H8···O5**′** and pyrimidine H6···O5**′** interactions.^9^ To implement the NBfix_0BPh_ correction in GROMACS, the corresponding Lennard–Jones parameters for the H5–OS and H4–OS atom-type pairs were added to the [nonbond_params] section of the GROMACS topology file. The H5–OS and H4–OS pairs represent the purine H8···O5**′** and pyrimidine H6···O5**′** contacts, respectively. The *σ_ij_* values were 0.25665010 and 0.26110460 nm for H5–OS and H4–OS, respectively, whereas *ε_ij_*was 0.211292 kJ mol**□**¹ for both pairs.

All simulations involving gHBfix corrections, including gHBfix21, gHBfix-18Ab, and gHBfix-18Ab*, were performed using PLUMED 2.8.4 to apply the additional biasing potentials.^30,31^ The parameterization of gHBfix-18Ab and benchmark simulations of the individual gHBfix parameter sets were performed using OL3 as the underlying force field. After selecting the final gHBfix-18Ab parameter sets, we constructed a composite model gHBfix-18Ab* by applying gHBfix-18Ab parameters and the NBfix_0BPh_ correction to the OL3_CP_ topology.^8,9^

### 2.3. Hydrogen-Bond Correction Parameters

The gHBfix corrections introduce additional pair-type-specific potentials acting on selected hydrogen–acceptor distances.^10,11^ For both the 6VA1 and 2KOC systems, donor-hydrogen/acceptor-atom pairs were restricted to local residue neighborhoods, as detailed in Text S3. These corrections do not directly modify atomic charges, bonded terms, or Lennard–Jones parameters. Instead, the correction strength depends on the donor and acceptor atom-group categories involved in each hydrogen-bond interaction.

For gHBfix21, correction strengths were taken directly from the original published parameter set without modification (Figure 3). This parameter set forms the 12-component correction matrix, which consists of two donor categories, NH and 2′-OH; and six acceptor categories, N, O, O4′, 2′-OH, bO, and nbO. The atomic groups included in each category are listed in Table S1.

**Figure 3.**
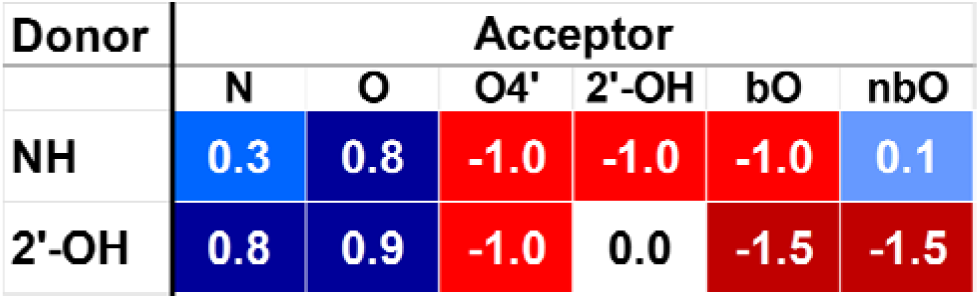
Published gHBfix21 correction matrix^10^. Numerical entries represent correction strengths as k_B_Tλ (kcal mol□¹). Blue and red cells indicate hydrogen-bond types favored and disfavored by the correction, respectively, with color intensity reflecting the magnitude of the correction strength.

Following the original gHBfix convention, λ is dimensionless, and the corresponding energy scale is given by k_B_Tλ. Therefore, in the figures, tables, and the main text, we report the correction strengths as k_B_Tλ in kcal mol□¹, where k_B_ is the Boltzmann constant and T is the simulation temperature. Positive and negative k_B_Tλ values favor and disfavor the corresponding hydrogen-bond interactions, respectively.

The gHBfix-18Ab correction used the same acceptor categories as those of gHBfix21 but separated the original NH donor category into NH and NHS donor categories. The resulting 18-component correction matrix consists of three donor categories, NH, NH□, and 2′-OH, and six acceptor categories (Figure 4). The initial NH□–N correction was assigned the most negative correction strength present in the gHBfix21 matrix. In the final matrix, the NH□–O and 2′-OH–N correction strengths were set to 0.0. The atomic groups belonging to each donor and acceptor category are listed in Table S2.

**Figure 4.**
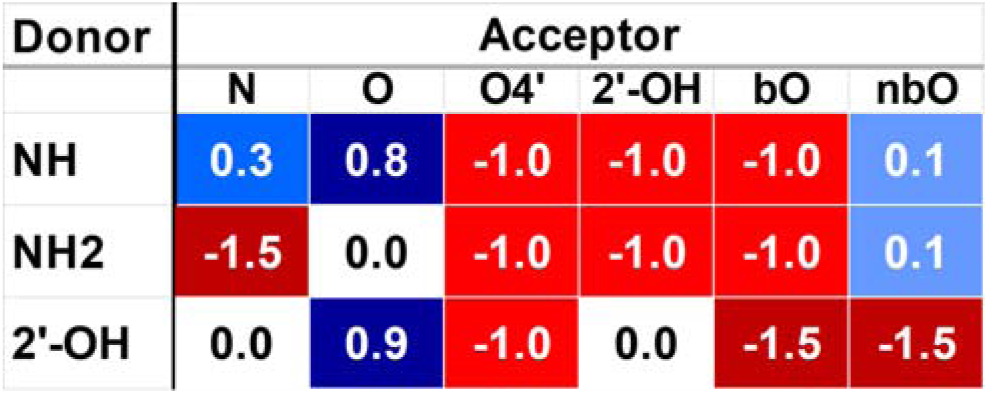
gHBfix-18Ab correction matrix developed in this study. The matrix separates the original NH donor category into the NH and NH□ donor categories. Numerical entries represent correction strengths reported as k_B_Tλ (kcal mol□¹). Blue and red cells indicate hydrogen-bond types favored and disfavored by the correction, respectively.

### 2.4. Molecular Dynamics Simulation Protocol

All MD simulations were performed using GROMACS 2022.5.^32^ Unless otherwise stated, the following MD protocol was used for all simulations. After energy minimization using the steepest-descent algorithm, each system was equilibrated for 125 ps in the constant-number, constant-volume, and constant-temperature (NVT) ensemble using a 1 fs timestep. Production simulations were performed in the constant-number, constant-pressure, and constant-temperature (NPT) ensemble at 300 K and 1 bar with a 2 fs timestep.

During production, temperature was controlled using a Nosé–Hoover thermostat, applied separately to the solute and solvent groups with a coupling time constant of 1.0 ps. Pressure was controlled isotropically using a Parrinello–Rahman barostat with a coupling time constant of 5.0 ps and a reference pressure of 1 bar. Electrostatic interactions were treated using the particle–mesh Ewald method with a real-space cutoff of 0.9 nm. Lennard–Jones interactions were truncated at 0.9 nm without a switching function, and long-range dispersion corrections were applied to the energy and pressure. Covalent bonds involving hydrogen atoms were constrained using the LINear Constraint Solver (LINCS) algorithm. Trajectory coordinates were saved every 50 000 steps (100 ps).

### 2.5. Accelerated Weight Histogram (AWH) Simulations of the 6VA1 A-Bulge System

The enhanced-sampling simulations were performed using the accelerated weight histogram (AWH) method implemented in GROMACS 2022.5.^12,13,32^ AWH is an adaptive-bias enhanced-sampling method that accelerates sampling along a predefined reaction coordinate while simultaneously estimating the corresponding free-energy profile. One-dimensional potentials of mean force (PMFs) were computed along the A-bulge pseudotorsion coordinate **θ**, which served as the reaction coordinate in the AWH calculations and was defined by the dihedral C16(C3**′**)–G7(C3**′**)–C5(C3**′**)–A6(N1) (Figure 5).^3^

**Figure 5.**
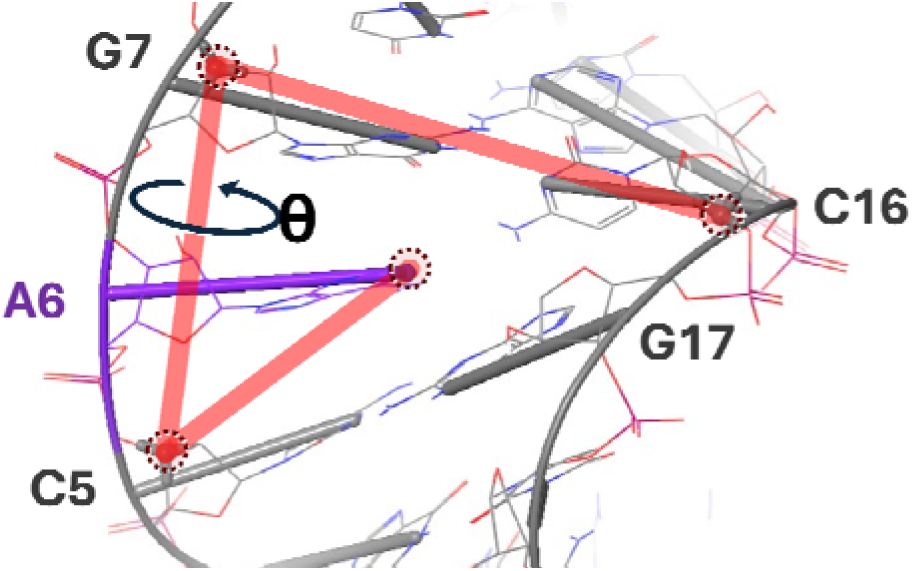
Definition of the A-bulge pseudotorsion coordinate (θ). The pseudotorsion coordinate θ was defined as the dihedral angle formed by C16(C3′), G7(C3′), C5(C3′), and A6(N1) in the A-bulge system. The first three atoms (C3′) are ribose atoms, whereas the fourth atom (N1) is the nucleobase atom of the bulged adenosine residue A6, shown in purple.

AWH simulations of the MAPT A-bulge system were performed for each force-field/correction setup under the conditions summarized in Table 2, including the simulation length and the number of independent runs. The reaction coordinate θ was implemented as a GROMACS dihedral pull coordinate and coupled to AWH through an external potential. The sampling range of θ was −100° to 100°. The bias force constant was 128 000 kJ mol□¹ rad□², and the diffusion constant was 5 × 10□□ rad² ps□¹. A local Boltzmann target distribution was employed with a target beta-scaling factor of 0.5, and the AWH growth mode was set to linear. AWH bias profiles were written every 50 000 steps (100 ps).

### 2.6. Reweighting of AWH-Biased Ensembles

Because AWH employs a time-dependent bias along the pseudotorsion coordinate θ, trajectory frames were reweighted to recover unbiased ensemble averages.^12,13^ All frames from the 0–1000 ns production interval were included in the analysis. For trajectory frame *i*, the correspondingdimensionless coordinate bias value reported in the GROMACS AWH output was denoted as *b_i_*. The frame weight was calculated as

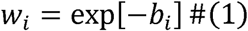

These weights were used in the reweighted NOE analysis described below.

### 2.7. NOE-Based Analysis of AWH Ensembles

Agreement with NMR-derived NOE distance restraints was evaluated using an indicator-based NOE sensitivity metric.^2^ For each NOE proton pair, the proton–proton distance was calculated for each analyzed trajectory frame using periodic boundary conditions. For a given proton pair with lower and upper bounds *L* and *U*, the indicator variable *h_i_* was defined as 1 when the distance *d_i_*satisfied *L* **≤** *d_i_* **≤** *U,* and as 0 otherwise. For AWH trajectories, the reweighted NOE sensitivity for that proton pair was calculated as

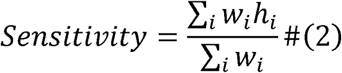

where the summation runs over all analyzed trajectory frames and *w_i_* is the AWH reweighting factor introduced above.

To evaluate local structural agreement around the A-bulge, 12 NOE proton pairs centered on A6 were selected from the NMR-derived NOE restraints (Figure 6), and this calculation was repeated for each pair. The mean NOE sensitivity was calculated as the arithmetic mean of the 12 pair-specific sensitivity values. The corresponding allowable distance ranges are summarized in Table S3.

**Figure 6.**
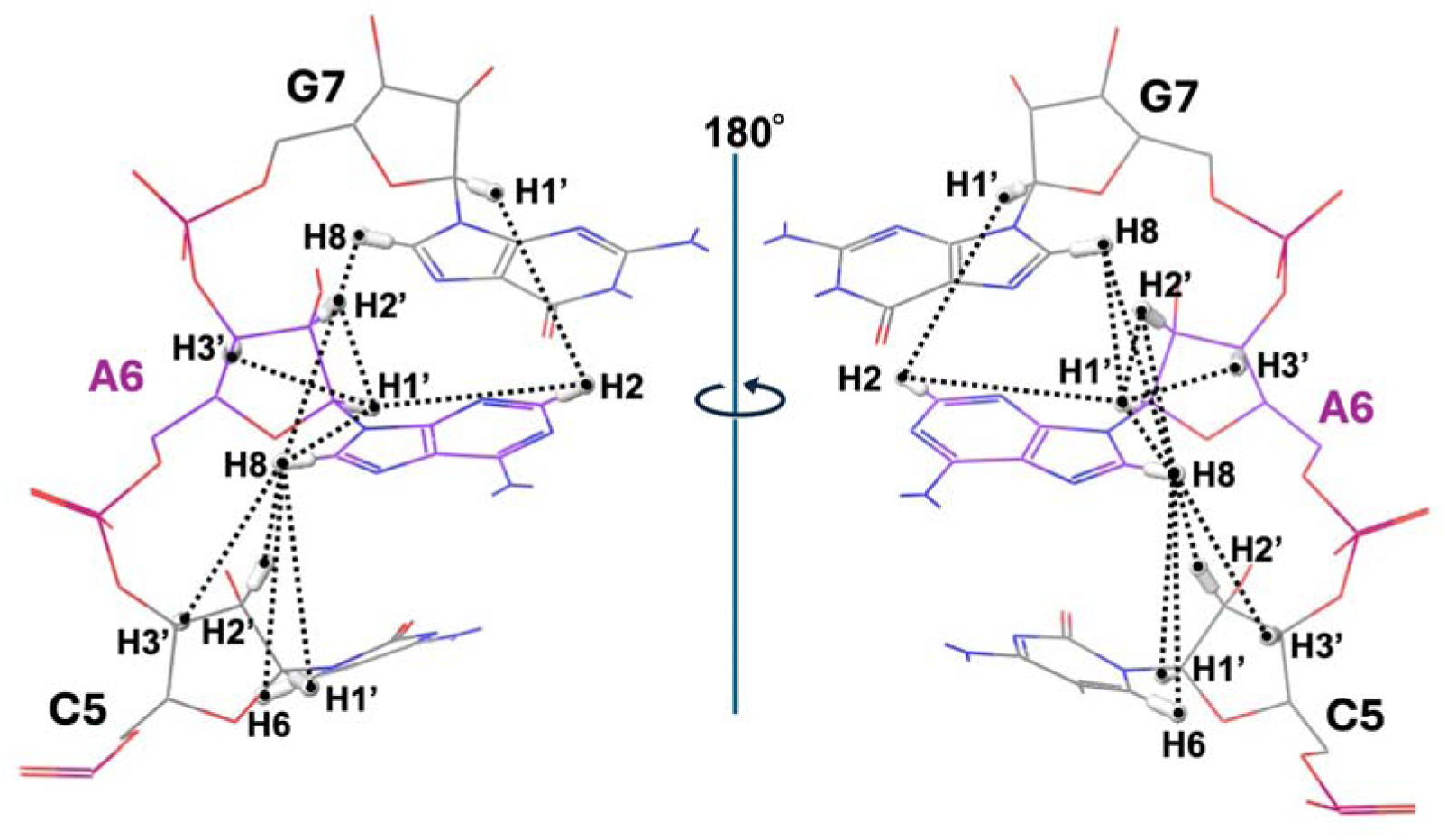
Twelve NOE proton pairs centered on residue A6 in the 6VA1 A-bulge structure. The selected 12 proton pairs, comprising intra-A6, A6–C5, and A6–G7 interactions, are shown in two views related by a 180° rotation, indicated by black dashed lines. NOE: nuclear Overhauser effect

### 2.8. Conventional MD Simulations of the cUUCGg Tetraloop

Conventional MD simulations of the cUUCGg tetraloop system were performed for the OL3 and gHBfix-18Ab* force-field models using the 2KOC NMR structure as the starting structure.^14^ For each force-field model, eight independent 4 μs trajectories were generated using different initial velocities, as summarized in Table 2. The simulations used the MD settings described above, except that no AWH bias was applied. For each trajectory, RNA all-atom root mean square deviation (RMSD) was calculated relative to the initial 2KOC NMR structure after least-squares fitting to the reference structure.

## 3. Results and Discussion

### 3.1. AWH-Derived Free-Energy Profile

As a representative example, we first present the one-dimensional PMF obtained using the AWH method^12,13^ with the standard OL3 force field^5–7^ along the pseudotorsion coordinate **θ** (Figure 5). The profile revealed two major free-energy minima corresponding to the NMR-supported stacked conformation and a non-native base-triple conformation (Figure 7). Consistent with previous MD studies,^3^ the OL3 force field also favored the non-native base-triple conformation in the AWH-derived PMF. To evaluate the convergence of the calculated PMF, profiles were calculated from cumulative trajectory lengths of 200–1000 ns and compared. The profiles obtained from the 800 and 1000 ns cumulative trajectories were nearly identical, indicating that the major features of the OL3 PMF, including the locations and relative free energies of the stacked and base-triple minima, were stable over the final part of the simulation (Figure 7). Therefore, the 1000 ns AWH-derived PMFs were used for the subsequent force-field comparisons.

**Figure 7.**
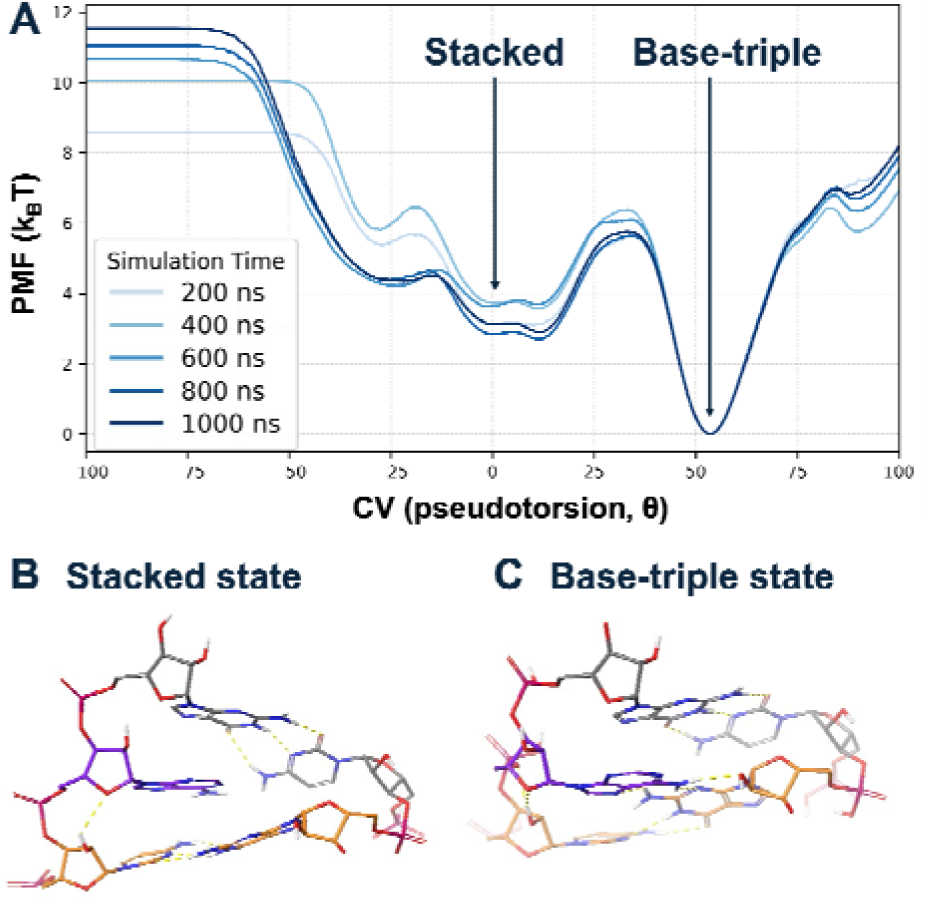
One-dimensional potentials of mean force (PMF) calculated using the AWH method along the pseudotorsion angle (θ) with the OL3 force field. (A) PMFs calculated from cumulative trajectory lengths of 200–1000 ns. Free energies are reported in units of k_B_T. Representative structures corresponding to the (B) stacked and (C) base-triple minima are shown.

### 3.2. Benchmarking of AMBER-Family RNA Force-Field Models and Correction Schemes

We next investigated whether other AMBER-family RNA force-field models and correction schemes improve the stacked/base-triple balance in the MAPT A-bulge. The reference OL3 force field^5–7^ was compared with six representative refinements and correction schemes: YIL,^15^ ROC,^17^ DE Shaw,^18^ CMAP,^19^ OL3_CP_,^8^ and gHBfix21^10^ (Table 2), covering torsional, phosphate, nonbonded, CMAP-based backbone, and hydrogen-bond refinement strategies.

As with OL3, all examined models and correction schemes except gHBfix21 favored the non-native base-triple state over the experimentally observed stacked state (Figure 8). gHBfix21 slightly favored the stacked state, although the free-energy difference between the two minima remained small. Thus, although gHBfix21 improved the stacked/base-triple balance relative to the other models, the base-triple state remained energetically comparable to the stacked state, rather than reproducing the NMR-supported dominance of the stacked state.

**Figure 8.**
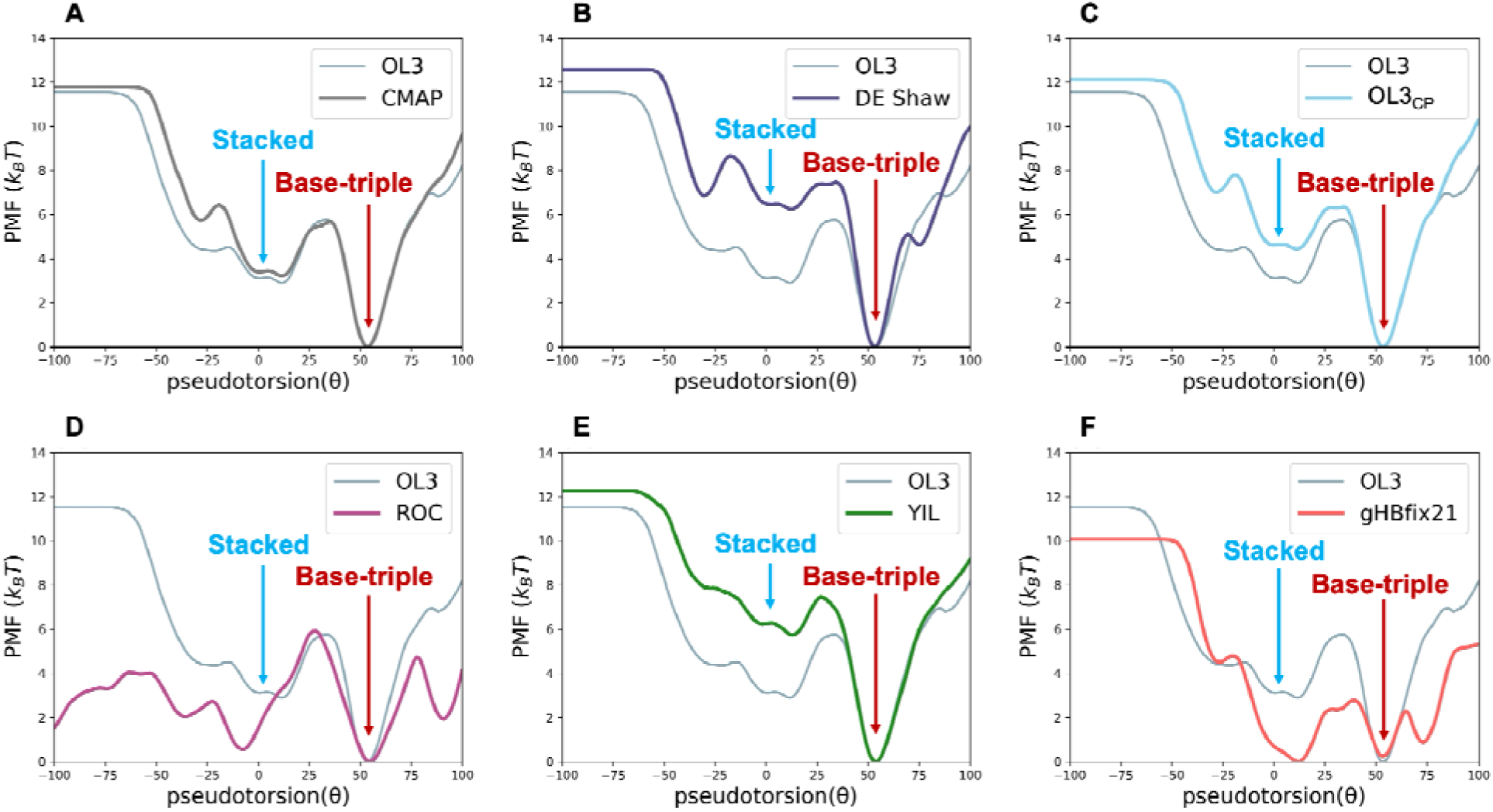
Comparison of one-dimensional PMFs among AMBER-family RNA force-field models and corrections. (A) CMAP, (B) DE Shaw, (C) OL3_CP_, (D) ROC, (E) YIL, and (F) gHBfix21. The OL3 PMF is shown in each panel for comparison. Free energies are reported in units of k_B_T.

To further assess whether the AWH ensembles are consistent with the experimental NMR data, we calculated reweighted NOE sensitivities for 12 A6-centered proton–proton distance restraints. These sensitivities quantify how well the local A6-centered conformational arrangement sampled by each force-field model satisfy the NMR-derived distance information and provide a structural diagnostic complementary to the one-dimensional PMF analysis.

All tested force-field models and correction schemes showed only moderate agreement with the A6-centered NOE distance ranges (Figure 9). The mean reweighted sensitivities ranged from approximately 0.57 to 0.67 across the benchmarked models. gHBfix21 showed the highest mean sensitivity in this set, consistent with its partial improvement in the stacked/base-triple balance, but the improvement was limited. Pair-specific A6-centered discrepancies remained, including low sensitivities for C5(H6)–A6(H8), A6(H8)–C5(H3**′**), and A6(H8)–G7(H8) (Table S4). Because these restraints involve distances between the A6 base and the adjacent C5 and G7 bases in the stacked state, their low sensitivities indicate that the stacked A6 arrangement was not fully reproduced.

**Figure 9.**
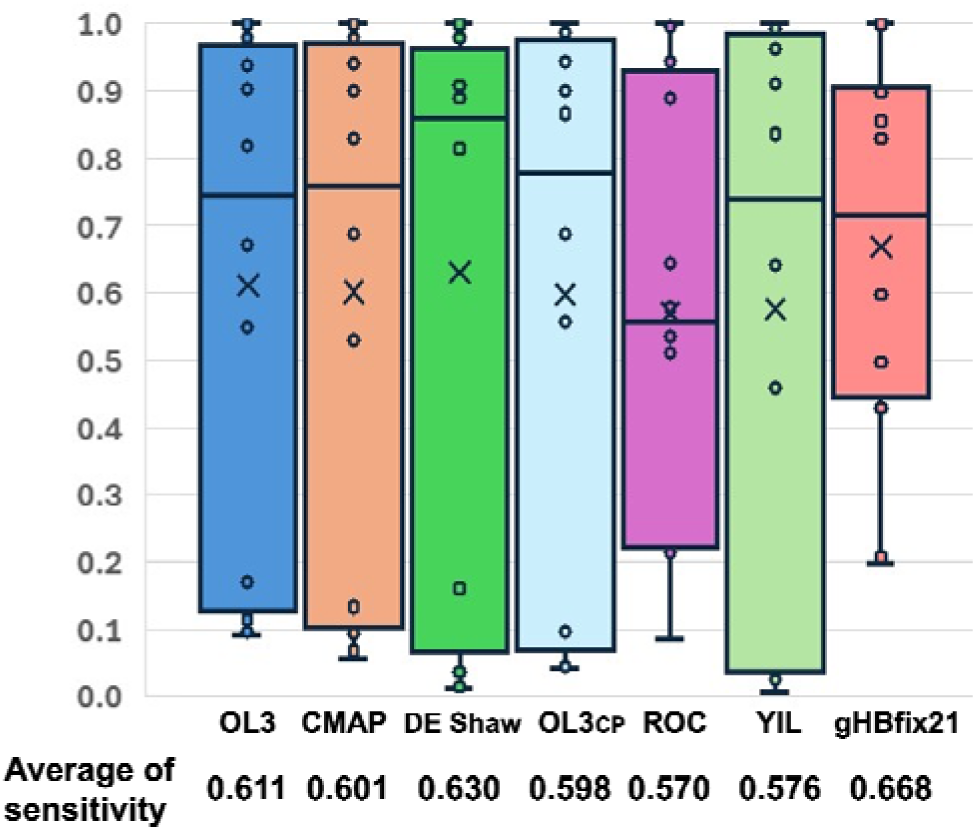
Comparison of reweighted NOE sensitivities across the tested RNA force fields. Box-and-whisker plots illustrate the distribution of sensitivities derived from the reweighted AWH ensembles. Crosses and the numerical values at the bottom indicate the mean sensitivity for each force field: OL3, CMAP, DE Shaw, OL3_CP_, ROC, YIL, and gHBfix21.

Taken together, these PMF and NOE analyses showed a consistent trend across the tested force-field models and corrections. All models tended to overestimate the stability of the non-native base-triple conformation relative to the experimentally observed stacked conformation. Although gHBfix21 improved this balance relative to other models and corrections, the stacked and base-triple minima remained close in free energy, and A6-centered NOE discrepancies persisted. These findings indicated that the conformational balance was still not satisfactorily reproduced, motivating the development of new force-field corrections aimed at more accurately reproducing the stacked A6 arrangement and its stability relative to the base-triple state.

### 3.3. Structural Basis for Targeting Base-Triple-Associated Hydrogen Bonds

To identify an appropriate force-field correction target, we first examined whether the conformational transition was associated primarily with backbone torsional changes. Backbone-dihedral analysis of the OL3 AWH trajectory (Text S1 and Figures S1 and S2) showed that only the ε and ζ torsions of C5 and A6 exhibited appreciable correlations with the pseudotorsion coordinate θ, whereas the remaining backbone dihedrals showed only weak correlations. Moreover, the ε/ζ changes between the stacked and base-triple states were continuous and relatively modest (approximately 10–17°), suggesting that large backbone torsional changes are unlikely to be the dominant feature distinguishing the stacked and base-triple conformations. Given the absence of large backbone torsional changes, we next examined local nonbonded interactions that differ between the two states and could directly contribute to their relative stability.

Specifically, we compared local interactions around A6 in the NMR stacked structure with those in a representative structure extracted from the base-triple minimum of the OL3 AWH-derived PMF (Figure 2). In the NMR stacked structure, the bulged A6 residue lies between the adjacent C5 and G7 bases and forms stacking interactions with both bases (Figure 2A). In the MD-derived base-triple structure, A6 rotates toward the adjacent C5–G17 Watson–Crick base pair. In this structure, two additional NH**□**-donor/N-acceptor hydrogen bonds are formed between A6 and G17: G17(N2–H)···A6(N7) and A6(N6–H)···G17(N3) (Figure 2B). These NH**□**-donor/N-acceptor hydrogen bonds therefore emerged as the primary correction target.

However, because the gHBfix21^10^ matrix assigns NH and NH**□** donor groups to the same NH-donor category (Figure 3), weakening the shared NH–N element to reduce these two NH**□**–N hydrogen bonds would inevitably also weaken other NH-donor/N-acceptor interactions, including the G17(N1–H)···C5(N3) hydrogen bond in the C5–G17 Watson–Crick base pair. We therefore introduced a separate NH**□** donor category in gHBfix-18Ab, allowing the NH**□**–N matrix element to be adjusted without changing the corresponding NH–N element. This donor-resolved classification formed the basis of the 18-component correction matrix used in the subsequent parameter refinement (Figure 4).

### 3.4. Design Rationale and Performance of gHBfix-18Ab

Using this donor-resolved classification, we developed the gHBfix-18Ab correction using the AWH-derived PMF and reweighted NOE sensitivities as benchmarks. Further details of the stepwise parameter refinement are provided in the Supporting Information (Text S2, Figures S3 and S4, and Table S5). Using the 18-component correction matrix, we first weakened the NH**□**–N correction to reduce the base-triple-associated hydrogen bonds (gHBfix-18Ab-pre1). In the resulting PMF, the base-triple-associated basin was raised relative to the stacked state, but an alternative low-free-energy state emerged, stabilized by an A6(N6–H)···C5(O2) hydrogen bond (Figures S3A and S5A). We therefore removed the NH**□**–O correction (gHBfix-18Ab-pre2), which suppressed this trapped state, but a second correction-induced state involving a U18(O2**′**–H)···A6(N1) hydrogen bond appeared (Figures S3B and S5B). In the final gHBfix-18Ab parameter set, the NH**□**–N correction was retained, whereas the NH**□**–O and 2**′**-OH–N correction terms were set to 0.0 (Figure S3C). This final parameter set made the stacked state the lowest-free-energy minimum, raised the base-triple-associated minimum, and eliminated the two correction-induced trapped basins (Figure 10A).

**Figure 10.**
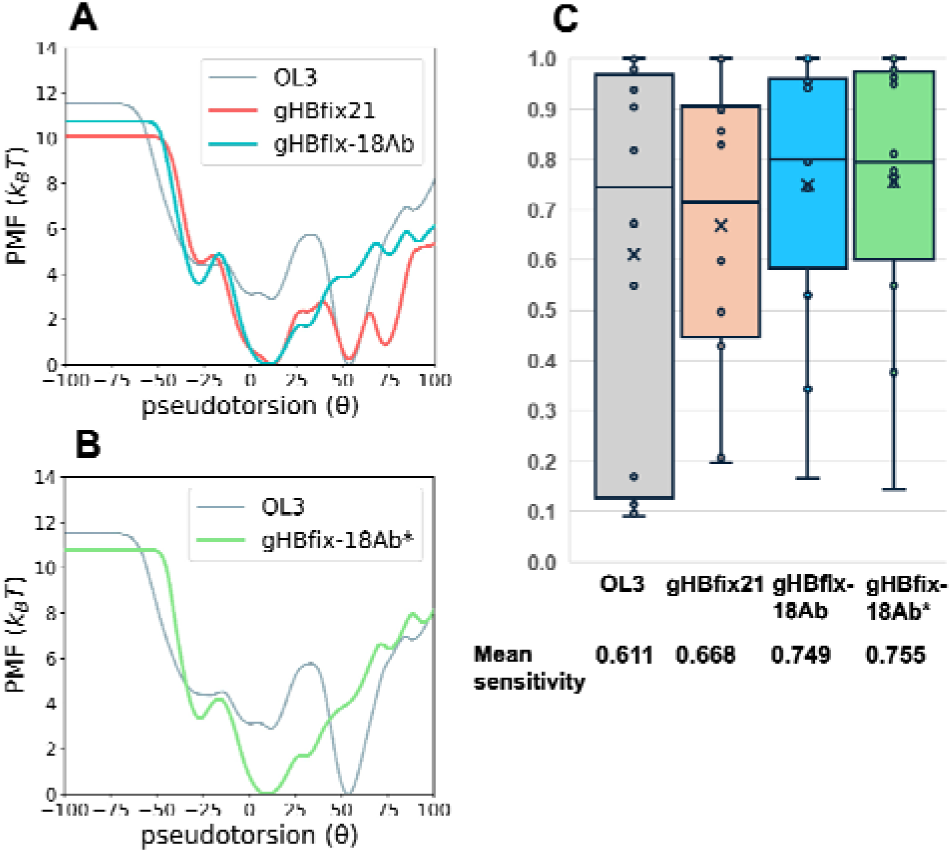
One-dimensional PMFs and reweighted NOE sensitivities obtained with OL3, gHBfix21, gHBfix-18Ab, and gHBfix-18Ab*. PMFs were computed from 1000 ns AWH simulations along the A-bulge pseudotorsion angle θ, with free energies reported in units of kBT. (A) Comparison of the PMFs obtained with OL3, gHBfix21, and gHBfix-18Ab. (B) Comparison of the PMFs obtained with OL3 and gHBfix-18Ab*. (C) Reweighted NOE sensitivities for OL3, gHBfix21, gHBfix-18Ab, and gHBfix-18Ab*. Box-and-whisker plots show the distributions of sensitivities for the 12 proton pairs around A6. Individual points represent pair-specific sensitivities, crosses indicate the mean values, and the numerical mean sensitivities are shown below the box plots.

After establishing the effect of gHBfix-18Ab in the standard OL3 force field, we constructed a composite force-field model, gHBfix-18Ab*, by combining gHBfix-18Ab with OL3_CP_ and NBfix_0BPh_. This composite setup follows previous recommendations in which gHBfix21 was used with a phosphate-corrected OL3 setup and NBfix_0BPh_ was subsequently added for simulations of small RNA motifs.^8–10^ The AWH-derived PMF of gHBfix-18Ab* retained the main features observed for gHBfix-18Ab in OL3, with the stacked state remaining the global minimum in the PMF and the base-triple-associated minimum raised relative to OL3 (Figure 10B).

The progressive improvements observed in the PMF were also reflected in the NOE sensitivity (Figure 10C and Figure S4). The mean reweighted NOE sensitivity increased from 0.611 with OL3 to 0.719–0.722 with the intermediate parameter sets (gHBfix-18Ab-pre1 and -pre2), to 0.749 with gHBfix-18Ab, and to 0.755 with gHBfix-18Ab*. Pair-resolved analysis of gHBfix-18Ab* showed higher sensitivities than OL3 for C5(H6)–A6(H8), A6(H8)–C5(H3**′**), and A6(H8)–G7(H8), although the improvement was smaller for A6(H8)–G7(H8) than for the two restraints involving C5 (Table S5). Together, the mean and pair-resolved NOE sensitivities indicate that the improvement over OL3 around A6 was retained in the composite model, gHBfix-18Ab*.

Because two restraints, A6(H8)–A6(H2**′**) and A6(H8)–G7(H8), still exhibited relatively low sensitivities in the gHBfix-18Ab* ensemble (0.144 and 0.376, respectively), we examined these proton pairs in more detail. For A6(H8)–A6(H2**′**), the reweighted mean proton–proton distance was 3.981 ± 0.216 Å, differing from the mean NMR distance (3.728 ± 0.073 Å) by only 0.253 Å (Figure 11). Moreover, the NOE upper bound (3.76 Å) lies very close to the mean NMR distance, and the NMR ensemble itself satisfies this restraint only partially (sensitivity = 0.600), with approximately 40% of the NMR structures exceeding the upper distance limit. For A6(H8)–G7(H8), the reweighted mean distance (4.872 ± 0.487 Å) differed from the mean NMR distance (5.095 ± 0.095 Å) by only 0.223 Å, whereas the NOE lower bound (5.00 Å) is likewise located immediately adjacent to the mean NMR value. Distances for the remaining proton pairs also showed close agreement between the gHBfix-18Ab* ensemble and the NMR structures, with ensemble-averaged distances generally differing by only a few tenths of an angstrom (Figure 11). Corresponding comparisons for the other force-field models are provided in Figure S6. Thus, the apparently low sensitivities of these two restraints primarily reflect the placement of the experimental NOE boundaries relative to the NMR distance distributions rather than substantial structural discrepancies between the reweighted ensemble and the NMR structures. Taken together, these analyses indicate that gHBfix-18Ab* reproduces the experimentally supported local A6 geometry despite the low sensitivities of these two individual restraints.

**Figure 11.**
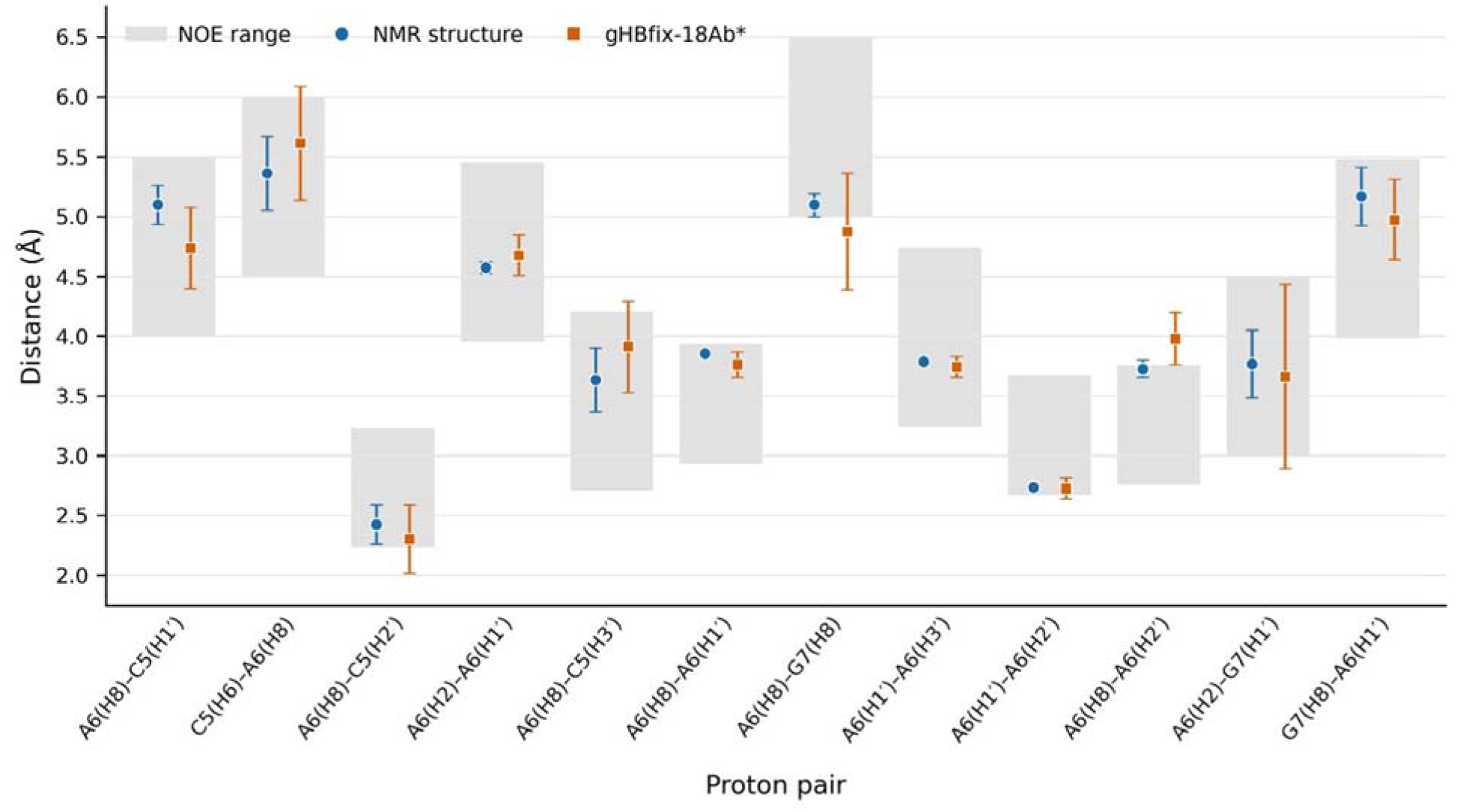
Comparison of A6-centered proton–proton distances in the 6VA1 NMR ensemble and the reweighted AWH ensemble obtained with gHBfix-18Ab*. Gray bands indicate the lower and upper bounds of the corresponding NOE distance restraints. Symbols and error bars represent the mean distance and population standard deviation, respectively. NMR values were calculated from the 20 deposited models of 6VA1, whereas the gHBfix-18Ab* values were calculated from the 1000 ns AWH trajectory after reweighting.

### 3.5. Reference-Structure Retention in the cUUCGg Tetraloop

To examine whether gHBfix-18Ab* causes marked structural destabilization outside the A-bulge, we evaluated retention of the reference structure with gHBfix-18Ab* using the cUUCGg tetraloop (PDB ID: 2KOC), a widely used RNA force-field benchmark. For OL3 and gHBfix-18Ab*, eight independent 4 **μ**s conventional MD trajectories were initiated from the NMR structure using different initial velocities.

With gHBfix-18Ab*, none of the eight trajectories showed a sustained increase in RNA all-atom RMSD relative to the initial NMR structure over 4 **μ**s (Figure 12). In the corresponding OL3 control simulations, several trajectories reached RMSD values of approximately 3–4 Å from the initial structure (Figure S7). Although gHBfix21 was not explicitly tested here, previous work reported stable cUUCGg tetraloop simulations with gHBfix21.^1^ Together, these results indicate that gHBfix-18Ab* did not produce a marked destabilizing effect in this cUUCGg tetraloop reference-structure retention test.

**Figure 12.**
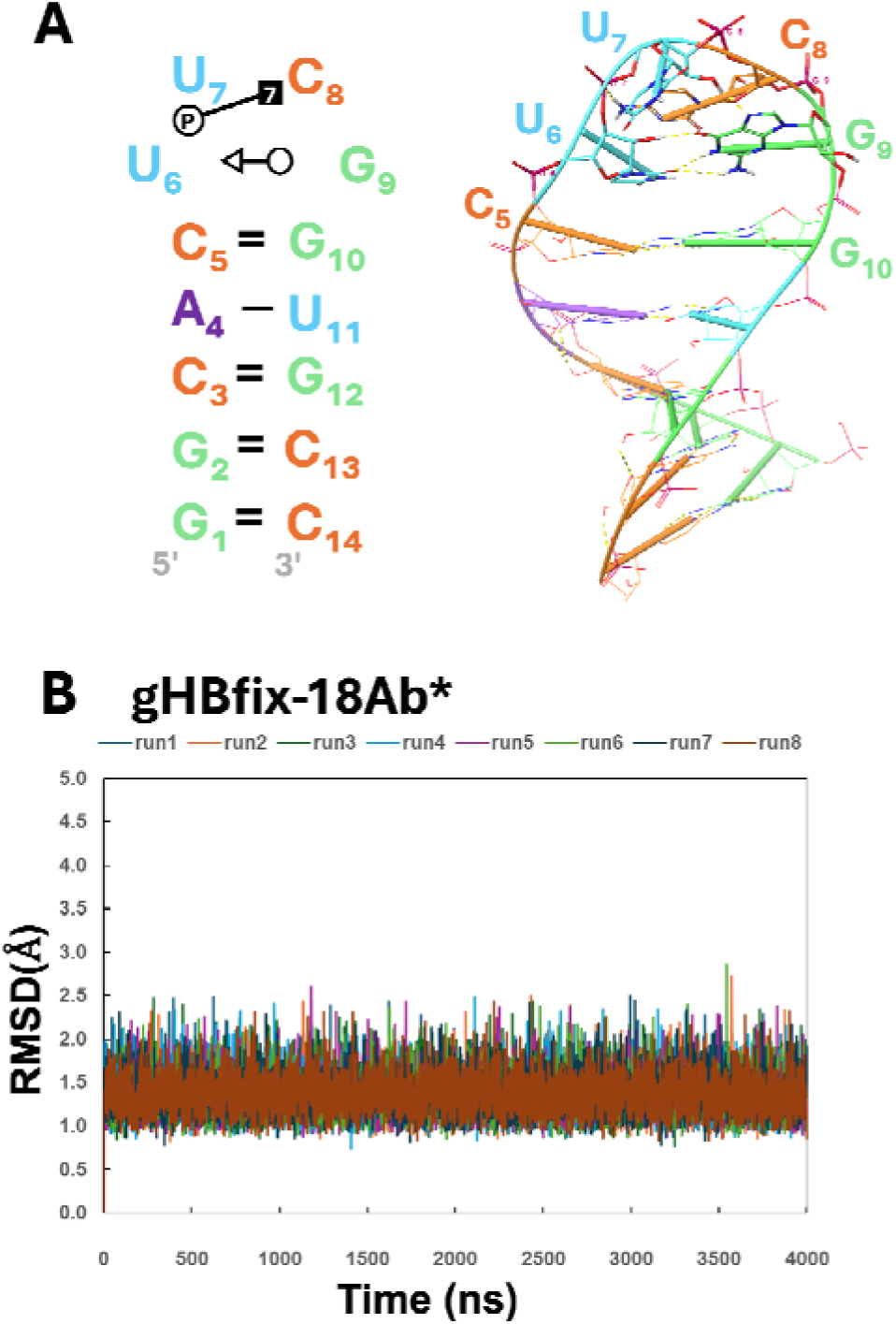
Sequence, NMR structure, and RNA all-atom root mean square deviation (RMSD) time series of the cUUCGg tetraloop with gHBfix-18Ab*. (A) Sequence and NMR structure of the cUUCGg tetraloop (PDB ID: 2KOC). (B) Time series of RNA all-atom RMSD relative to the initial NMR structure for gHBfix-18Ab*, calculated from eight independent 4 μs conventional MD trajectories initiated with different initial velocities.

### 3.6. Limitations and Future Directions

The A-bulge benchmark was limited to a single MAPT sequence context (5′-GCAGU/5′-ACGU), and the evaluation of non-A-bulge motifs was focused only on reference-structure retention in the cUUCGg tetraloop. Further evaluation of A-bulge motifs in different sequence contexts and additional RNA motifs will be needed to assess the broader applicability of gHBfix-18Ab*. Evaluating whether the improved description of A-bulge conformational ensembles translates into more accurate modeling of RNA–ligand interactions will be important, including ligand binding modes and binding energetics.

## 4. Conclusions

Using the MAPT A-bulge benchmark, we identified an imbalance between stacked and base-triple states as a remaining problem in the tested AMBER-family RNA force-field models. Comparison of the stacked and base-triple conformations revealed that the imbalance primarily arose from overly favorable NH**□**–N hydrogen bonds between A6 and the adjacent C5–G17 Watson–Crick base pair, indicating the need to refine NH**□**–N interactions independently of the corresponding NH–N interactions. Guided by this finding, we developed gHBfix-18Ab, which distinguishes NH and NH**□** donor categories, allowing selective adjustment of the base-triple-associated NH**□**–N interactions without altering the corresponding NH–N interactions. We further combined gHBfix-18Ab with the previously developed OL3_CP_ phosphate Lennard–Jones correction and NBfix_0BPh_ pair-specific Lennard–Jones correction to generate the composite force-field model gHBfix-18Ab*. The resulting gHBfix-18Ab* model restored the stacked state as the global minimum in the AWH-derived PMF and improved agreement with experimental NOE data around A6. To assess whether gHBfix-18Ab* destabilizes a non-A-bulge RNA motif, we performed a reference-structure retention test using the cUUCGg tetraloop, a well-established RNA force-field benchmark. No marked destabilization of the cUUCGg tetraloop was observed with gHBfix-18Ab*. Together, these findings demonstrate that targeted refinement of hydrogen-bond interactions provides a practical strategy for systematic improvement of AMBER-family RNA force fields toward more accurate modeling of noncanonical RNA motifs.

## ASSOCIATED CONTENT

### Supporting Information

Backbone-dihedral analysis and A-bulge state definitions; stepwise refinement of the gHBfix-18Ab parameter set and associated structural artifacts; donor–acceptor pair-selection rules for gHBfix calculations; atomic-group definitions for gHBfix21 and gHBfix-18Ab; NMR-derived NOE distance restraints, reweighted NOE sensitivities, and proton–proton distance comparisons; and RNA all-atom RMSD time series for the cUUCGg tetraloop (PDF).

### Data and Software Availability

#### Data

The gHBfix-18Ab parameters for the RNA A-bulge motif are available at https://github.com/IkeguchiLab/a-bulge-ff. The molecular dynamics trajectories and input files generated in this study have been deposited into the Biological Structure Model Archive (BSM-Arc) at Protein Data Bank Japan (PDBj) under accession number BSM00112.

#### Software

Molecular dynamics simulations were performed using GROMACS 2022.5 together with the PLUMED 2.8.4 plugin. Both software packages are open source and are publicly available at https://www.gromacs.org and https://www.plumed.org, respectively.

## AUTHOR INFORMATION

### Author Contributions

M.I. conceived and designed the study. T.K. performed the calculations and primarily carried out the analyses. T.E. and T.Y. contributed to the analyses and discussions. The manuscript was written through the contributions of all authors. All authors have given approval for the final version of the manuscript.

## ACKNOWLEDGMENT

This research used the computational resources at Yokohama City University, Tsurumi Campus, Japan. The authors acknowledge the use of Gemini (Google) and ChatGPT (OpenAI) for language editing, as well as for assistance in generating Python scripts used for data analysis and visualization. All AI-assisted scripts, analyses, and manuscript text were independently reviewed and verified by the authors, who take full responsibility for the content and scientific integrity of this work.

## ABBREVIATIONS

AWH: accelerated weight histogram
MAPT: microtubule-associated protein tau
MD: molecular dynamics
NMR: nuclear magnetic resonance
NOE: nuclear Overhauser effect
PDB: Protein Data Bank
PMF: potential of mean force
RMSD: root mean square deviation

