## Supporting Information for "Development of force-field corrections for the RNA A-bulge motif"

**Table of Contents**

Text S1. Backbone-Dihedral Analysis around the A-bulge. ..... S2
Text S2. Stepwise refinement of the gHBfix-18Ab parameter set. ..... S2
Text S3. Selection of donor–acceptor pairs for gHBfix calculations. ..... S2
Table S1. Atomic-group types corrected by gHBfix21. ..... S3
Table S2. Atomic-group types corrected by gHBfix-18Ab. ..... S3
Table S3. NOE distances and allowable ranges used for sensitivity calculations. ..... S4
Table S4. Reweighted NOE sensitivities for contemporary RNA force fields. ..... S4
Table S5. Reweighted NOE sensitivities during the stepwise gHBfix parameterization. ..... S5
Figure S1. Definitions of backbone torsions and A-bulge states. ..... S6
Figure S2. Correlation between backbone dihedral angles and pseudotorsion angle θ. ..... S7
Figure S3. Stepwise refinement of the gHBfix-18Ab correction parameters for the MAPT A-bulge .. S8
Figure S4. Box-and-whisker plots and mean values of the reweighted NOE sensitivities obtained from MD trajectories using each gHBfix parameter set. ..... S9
Figure S5. Structural artifacts observed with gHBfix-18Ab-pre1 and gHBfix-18Ab-pre2. ... S10
Figure S6. Comparison of proton–proton distances around the A6 bulge across the NMR ensemble and AWH simulations. ... S11
Figure S7. RNA all-atom RMSD time series of the cUUCGg tetraloop with OL3. ... S12

**Text S1.** Backbone-dihedral analysis around the A-bulge

To characterize backbone changes associated with the A-bulge pseudotorsion coordinate θ, we analyzed the α, β, γ, δ, ε, and ζ backbone dihedral angles of residues surrounding the A-bulge in the OL3 AWH trajectory (Figures S1 and S2). Trajectory frames were classified into the major-groove, stacked, base-triple, and unstacked regions according to θ, as shown in Figure S1B.

Figure S2 shows the relationships between θ and the individual backbone dihedral angles. Among the analyzed torsions, the ε and ζ dihedrals of C5 and A6 exhibited the strongest correlations with θ, whereas the remaining backbone dihedrals showed comparatively weak correlations. Pearson correlation coefficients are summarized in Figure S2. The absolute differences in the mean dihedral angles between the stacked and base-triple states were 14.7° and 11.9° for the ε and ζ torsions of C5, respectively, and 9.6° and 16.6° for those of A6, respectively.

**Text S2.** Stepwise refinement of the gHBfix-18Ab parameter set

The gHBfix-18Ab parameter set was refined stepwise by adjusting the $NH_2$–N, $NH_2$–O, and 2′-OH–N correction terms and comparing the resulting AWH-derived PMFs and reweighted A6-centered NOE sensitivities (Figures S3 and S4).

In gHBfix-18Ab-pre1, the $NH_2$–N correction was applied to weaken the base-triple-associated hydrogen bonds (Figure S3A). This correction raised the base-triple-associated basin relative to the stacked state and introduced an alternative low-free-energy state stabilized by an A6(N6–H)···C5(O2) hydrogen bond (Figures S3A and S5A).

In gHBfix-18Ab-pre2, the $NH_2$–O correction term was set to 0.0 while retaining the $NH_2$–N correction (Figure S3B). This modification suppressed the alternative state stabilized by A6(N6–H)···C5(O2). Inspection of the resulting PMF, together with a representative structure extracted from the new minimum, revealed a second alternative state involving a U18(O2′–H)···A6(N1) hydrogen bond (Figures S3B and S5B).

**Text S3.** Selection of donor–acceptor pairs for gHBfix calculations

To restrict gHBfix corrections to local interactions, donor–acceptor pairs were generated from overlapping residue neighborhoods. For 6VA1, using Figure 1 numbering, the ranges on the two strands were 1–3/19–21, 2–4/18–20, 3–6/16–19, 5–7/16–17, 6–9/14–17, 8–10/13–15, and 9–11/12–14. For 2KOC, using Figure 12A numbering, the neighborhoods were 1–3/12–14, 2–4/11–13, 3–5/10–12, 4–6/9–11, and 5–10. Within each neighborhood, all donor-hydrogen/acceptor-atom combinations defined for the corresponding gHBfix parameter set (Tables S1 and S2) were included, with duplicate atom pairs from overlapping neighborhoods retained only once.

**Table S1.** List of eight atomic-group types involved in RNA nucleotide interactions corrected by gHBfix21

| Donors | A | C | G | U |
|---|---|---|---|---|
| NH(base) | N6(H61, H62) | N4(H41, H42) | N1(H1),<br>N2(H21, H22) | N3(H3) |
| 2′-OH(sugar) | O2′(HO2′) | O2′(HO2′) | O2′(HO2′) | O2′(HO2′) |
| **Acceptors** | **A** | **C** | **G** | **U** |
| N(base) | N1, N3, N7 | N3 | N3, N7 | - |
| O(base) | - | O2 | O6 | O2, O4 |
| O(sugar) | O4′ | O4′ | O4′ | O4′ |
| 2′-OH(sugar) | O2′ | O2′ | O2′ | O2′ |
| bO(phosphate) | O3′, O5′ | O3′, O5′ | O3′, O5′ | O3′, O5′ |
| nbO(phosphate) | pro-RP, pro-SP | pro-RP, pro-SP | pro-RP, pro-SP | pro-RP, pro-SP |

**Table S2.** List of nine atomic-group types involved in RNA nucleotide interactions corrected by gHBfix-18Ab

| Donors | A | C | G | U |
|---|---|---|---|---|
| NH(base) | - | - | N1(H1) | N3(H3) |
| $NH_2$(base) | N6(H61, H62) | N4(H41, H42) | N2(H21, H22) | - |
| 2′-OH(sugar) | O2′(HO2′) | O2′(HO2′) | O2′(HO2′) | O2′(HO2′) |
| **Acceptors** | **A** | **C** | **G** | **U** |
| N(base) | N1, N3, N7 | N3 | N3, N7 | - |
| O(base) | - | O2 | O6 | O2, O4 |
| O(sugar) | O4′ | O4′ | O4′ | O4′ |
| 2′-OH(sugar) | O2′ | O2′ | O2′ | O2′ |
| bO(phosphate) | O3′, O5′ | O3′, O5′ | O3′, O5′ | O3′, O5′ |
| nbO(phosphate) | pro-RP, pro-SP | pro-RP, pro-SP | pro-RP, pro-SP | pro-RP, pro-SP |

**Table S3.** NOE distances and allowable ranges used for sensitivity calculations (PDB ID: 6VA1)
Twelve NOE distance restraints around the A6 residue within the A-bulge region are listed along with their uncertainty ranges.

| Proton Pair | Lower bound (Å) | Upper bound (Å) |
|---|---|---|
| A6(H8)-C5(H1′) | 4.00 | 5.50 |
| C5(H6)-A6(H8) | 4.50 | 6.00 |
| A6(H8)-C5(H2′) | 2.23 | 3.23 |
| A6(H2)-A6(H1′) | 3.95 | 5.45 |
| A6(H8)-C5(H3′) | 2.71 | 4.21 |
| A6(H8)-A6(H1′) | 2.93 | 3.93 |
| A6(H8)-G7(H8) | 5.00 | 6.50 |
| A6(H1′)-A6(H3′) | 3.24 | 4.74 |
| A6(H1′)-A6(H2′) | 2.67 | 3.67 |
| A6(H8)-A6(H2′) | 2.76 | 3.76 |
| A6(H2)-G7(H1′) | 3.00 | 4.50 |
| G7(H8)-A6(H1′) | 3.98 | 5.48 |

**Table S4.** Reweighted NOE sensitivities for contemporary RNA force fields
Values represent the reweighted sensitivity for each of the 12 proton pairs around the A6 residue, calculated from AWH-MD trajectories. These data correspond to the results shown in Figure 9. The sensitivities were computed as weighted averages based on the bias potential applied during sampling. Experimental NOE distance restraints and proton pair identifiers are based on PDB ID: 6VA1.

| Proton Pair | OL3 | CMAP | DE Shaw | $OL3_{CP}$ | ROC | YIL | gHBfix21 |
|---|---|---|---|---|---|---|---|
| A6(H8)-C5(H1′) | 0.549 | 0.530 | 0.829 | 0.557 | 0.510 | 0.459 | 0.855 |
| C5(H6)-A6(H8) | 0.169 | 0.133 | 0.036 | 0.096 | 0.223 | 0.040 | 0.496 |
| A6(H8)-C5(H2′) | 0.818 | 0.829 | 0.909 | 0.866 | 0.577 | 0.836 | 0.598 |
| A6(H2)-A6(H1′) | 0.999 | 0.999 | 1.000 | 0.999 | 0.996 | 0.998 | 1.000 |
| A6(H8)-C5(H3′) | 0.114 | 0.068 | 0.015 | 0.044 | 0.213 | 0.007 | 0.429 |
| A6(H8)-A6(H1′) | 0.979 | 0.980 | 0.978 | 0.987 | 0.944 | 0.993 | 0.907 |
| A6(H8)-G7(H8) | 0.091 | 0.057 | 0.013 | 0.041 | 0.220 | 0.025 | 0.206 |
| A6(H1′)-A6(H3′) | 1.000 | 1.000 | 1.000 | 1.000 | 1.000 | 1.000 | 1.000 |
| A6(H1′)-A6(H2′) | 0.904 | 0.901 | 0.888 | 0.899 | 0.890 | 0.911 | 0.829 |
| A6(H8)-A6(H2′) | 0.096 | 0.093 | 0.160 | 0.060 | 0.084 | 0.035 | 0.198 |
| A6(H2)-G7(H1′) | 0.672 | 0.687 | 0.814 | 0.687 | 0.536 | 0.641 | 0.600 |
| G7(H8)-A6(H1′) | 0.938 | 0.941 | 0.916 | 0.943 | 0.644 | 0.963 | 0.897 |
| Average of sensitivity | 0.611 | 0.601 | 0.630 | 0.598 | 0.570 | 0.576 | 0.668 |

**Table S5.** Reweighted NOE sensitivities during the stepwise gHBfix parameterization

Values represent the reweighted sensitivity for each of the 12 proton pairs around the A6 residue for the baseline OL3 force field and the intermediate correction sets (gHBfix-18Ab-pre1, gHBfix-18Ab-pre2, gHBfix-18Ab, gHBfix-18Ab*). These data correspond to the results shown in Figure S4. Calculation methods and experimental reference data are identical to those described in Table S4.

| Proton Pair | OL3 | gHBfix-18Ab-pre1 | gHBfix-18Ab-pre2 | gHBfix-18Ab | gHBfix-18Ab* |
|---|---|---|---|---|---|
| A6(H8)-C5(H1′) | 0.549 | 0.927 | 0.919 | 0.962 | 0.978 |
| C5(H6)-A6(H8) | 0.169 | 0.716 | 0.746 | 0.805 | 0.777 |
| A6(H8)-C5(H2′) | 0.818 | 0.518 | 0.509 | 0.529 | 0.549 |
| A6(H2)-A6(H1′) | 0.999 | 0.989 | 0.963 | 1.000 | 1.000 |
| A6(H8)-C5(H3′) | 0.114 | 0.672 | 0.693 | 0.744 | 0.811 |
| A6(H8)-A6(H1′) | 0.979 | 0.908 | 0.912 | 0.955 | 0.950 |
| A6(H8)-G7(H8) | 0.091 | 0.263 | 0.307 | 0.342 | 0.376 |
| A6(H1′)-A6(H3′) | 1.000 | 1.000 | 1.000 | 1.000 | 1.000 |
| A6(H1′)-A6(H2′) | 0.904 | 0.788 | 0.808 | 0.795 | 0.752 |
| A6(H8)-A6(H2′) | 0.096 | 0.234 | 0.171 | 0.165 | 0.144 |
| A6(H2)-G7(H1′) | 0.672 | 0.693 | 0.719 | 0.755 | 0.765 |
| G7(H8)-A6(H1′) | 0.938 | 0.926 | 0.913 | 0.942 | 0.963 |
| Average of sensitivity | 0.611 | 0.719 | 0.722 | 0.749 | 0.755 |

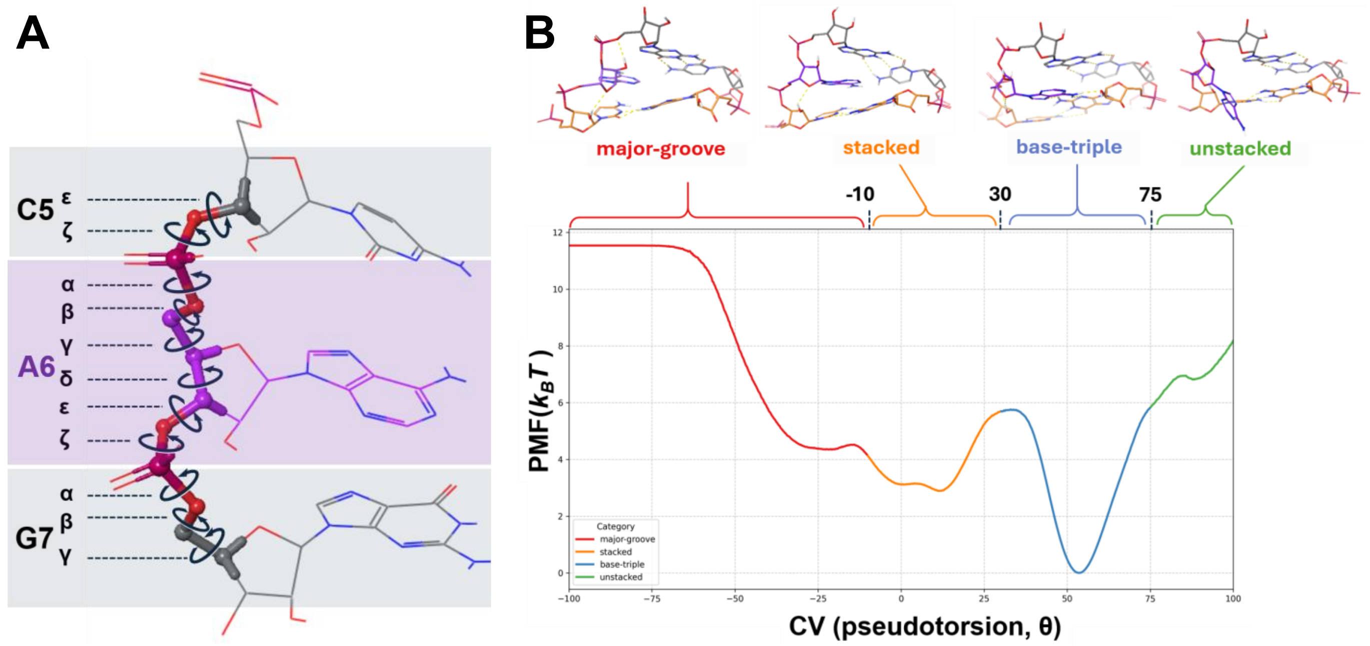


**Figure S1.** Definitions related to the analysis of backbone torsions and A-bulge states based on the one-dimensional PMF

(A) Backbone dihedral angles of residues surrounding the A-bulge. Each dihedral angle is defined by the following four atoms, where $\boldsymbol{i}$ represents the current nucleotide, $\boldsymbol{i-1}$ is the 5′-side neighbor, and $\boldsymbol{i+1}$ is the 3′-side neighbor: α($O3'_{i-1}$–$P_i$–$O5'_i$–$C5'_i$), β($P_i$–$O5'_i$–$C5'_i$–$C4'_i$), γ($O5'_i$–$C5'_i$–$C4'_i$–$C3'_i$), δ($C5'_i$–$C4'_i$–$C3'_i$–$O3'_i$), ε($C4'_i$–$C3'_i$–$O3'_i$–$P_{i+1}$), ζ($C3'_i$–$O3'_i$–$P_{i+1}$–$O5'_{i+1}$).

(B) Definition of the A-bulge states based on the 1000 ns one-dimensional PMF (OL3 force field) using the pseudotorsion angle (θ). θ ranges were classified as follows: -100° to -10° (major-groove; red), -10° to 30° (stacked; yellow), 30° to 75° (base-triple; blue), 75° to 100° (unstacked; green).

### Dihedral Angle Analysis for C5

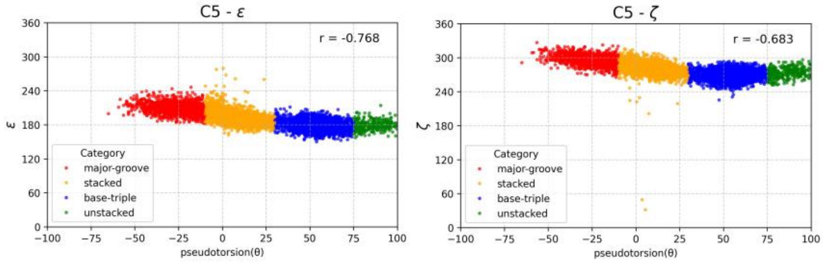


### Dihedral Angle Analysis for A6

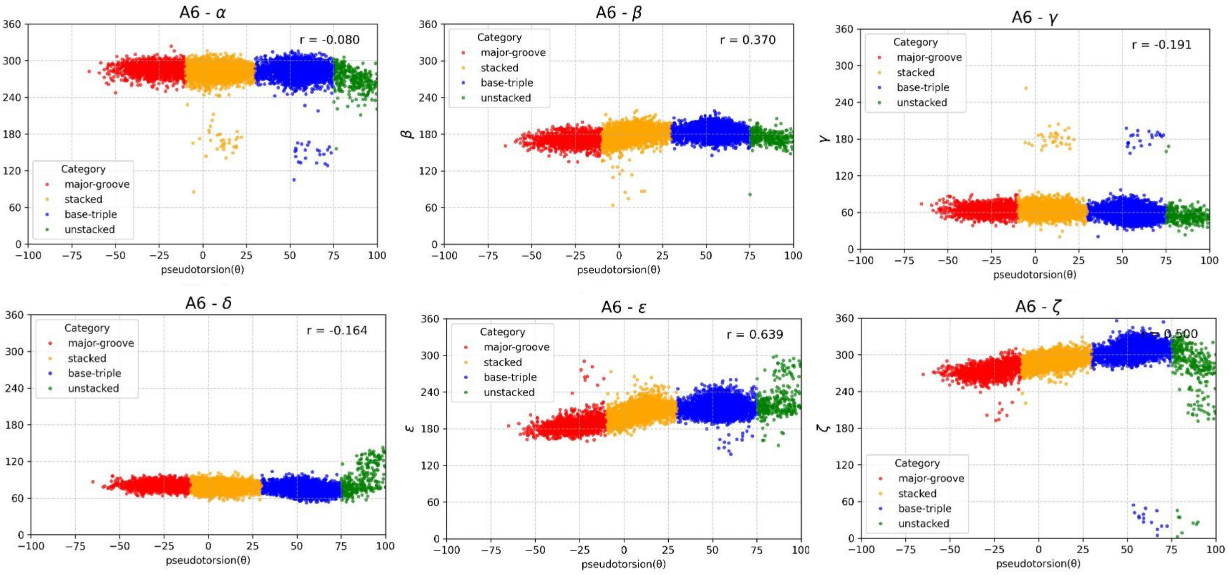


### Dihedral Angle Analysis for G7

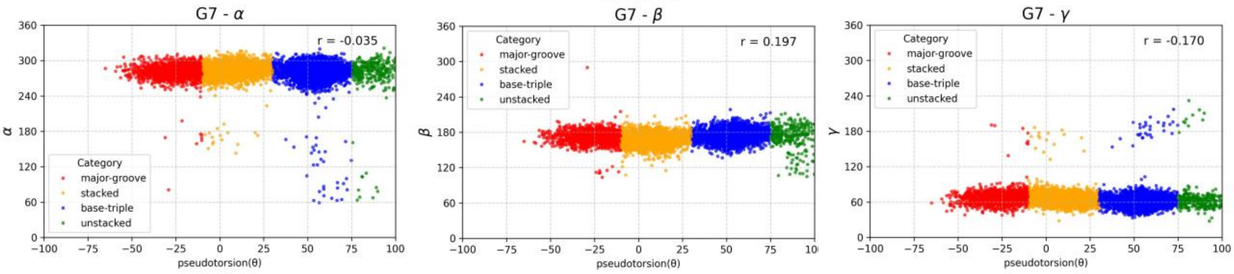


**Figure S2.** Correlation between backbone dihedral angles of residues near the A-bulge (C5, A6, G7) and the pseudotorsion angle (θ), calculated from the AWH MD trajectory (OL3 force field, 1000 ns, 10,001 frames) Each panel plots a backbone dihedral angle on the y-axis versus the pseudotorsion angle (θ) on the x-axis, along with the corresponding correlation coefficient.

θ ranges were classified as follows: -100° to -10° (major-groove; red), -10° to 30° (stacked; yellow), 30° to 75° (base-triple; blue), 75° to 100° (unstacked; green).

**A gHBfix-18Ab-pre1**

| Donor | Acceptor | | | | | |
|---|---|---|---|---|---|---|
| | N | O | O4' | 2'-OH | bO | nbO |
| NH | 0.3 | 0.8 | -1.0 | -1.0 | -1.0 | 0.1 |
| NH2 | -1.5 | 0.8 | -1.0 | -1.0 | -1.0 | 0.1 |
| 2'-OH | 0.8 | 0.9 | -1.0 | 0.0 | -1.5 | -1.5 |

**B gHBfix-18Ab-pre2**

| Donor | Acceptor | | | | | |
|---|---|---|---|---|---|---|
| | N | O | O4' | 2'-OH | bO | nbO |
| NH | 0.3 | 0.8 | -1.0 | -1.0 | -1.0 | 0.1 |
| NH2 | -1.5 | 0.0 | -1.0 | -1.0 | -1.0 | 0.1 |
| 2'-OH | 0.8 | 0.9 | -1.0 | 0.0 | -1.5 | -1.5 |

**C gHBfix-18Ab**

| Donor | Acceptor | | | | | |
|---|---|---|---|---|---|---|
| | N | O | O4' | 2'-OH | bO | nbO |
| NH | 0.3 | 0.8 | -1.0 | -1.0 | -1.0 | 0.1 |
| NH2 | -1.5 | 0.0 | -1.0 | -1.0 | -1.0 | 0.1 |
| 2'-OH | 0.0 | 0.9 | -1.0 | 0.0 | -1.5 | -1.5 |

**D gHBfix-18Ab***

| Donor | Acceptor | | | | | |
|---|---|---|---|---|---|---|
| | N | O | O4' | 2'-OH | bO | nbO |
| NH | 0.3 | 0.8 | -1.0 | -1.0 | -1.0 | 0.1 |
| NH2 | -1.5 | 0.0 | -1.0 | -1.0 | -1.0 | 0.1 |
| 2'-OH | 0.0 | 0.9 | -1.0 | 0.0 | -1.5 | -1.5 |

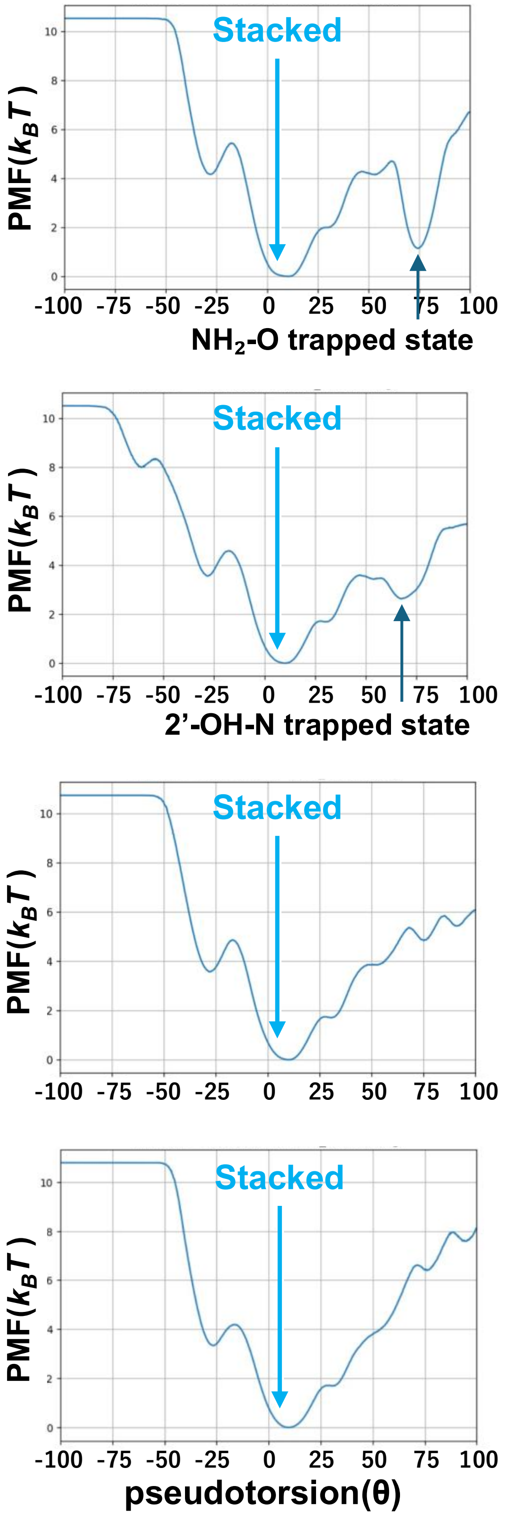


**Figure S3.** Stepwise refinement of the gHBfix-18Ab correction parameters for the MAPT A-bulge

For each parameter set, the left panel shows the hydrogen-bond correction matrix and the right panel shows the corresponding one-dimensional PMF along the A-bulge pseudotorsion coordinate θ. Correction-matrix entries are reported as kBTλ in kcal mol$^{-1}$. Blue and red cells indicate hydrogen-bond types favored and disfavored by the correction, respectively. The parameter sets shown are (A) gHBfix-18Ab-pre1, (B) gHBfix-18Ab-pre2, (C) gHBfix-18Ab, and (D) gHBfix-18Ab*.

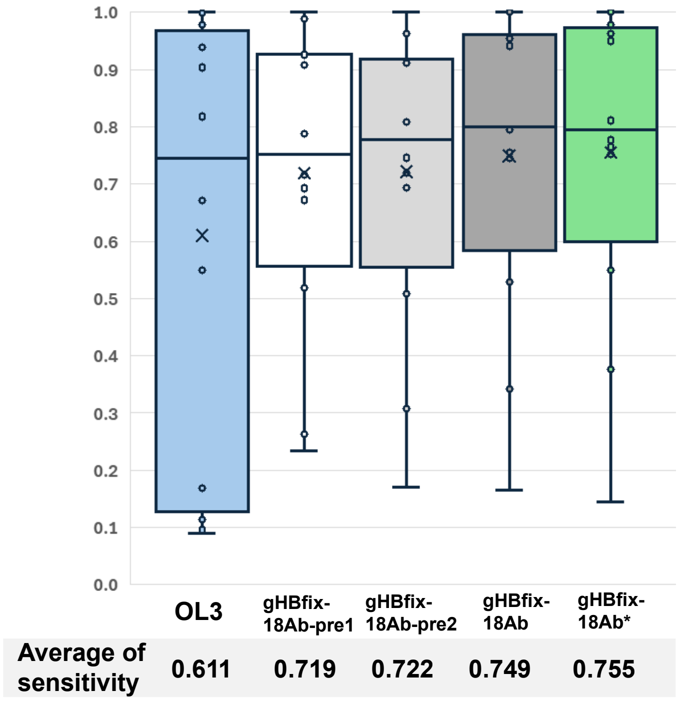


**Figure S4.** Box-and-whisker plots and mean values of the reweighted NOE sensitivities obtained from MD trajectories using each gHBfix parameter set

Box-and-whisker plots represent the distribution of sensitivities calculated from MD trajectories. Mean values are provided below the respective x-axis labels. The composition and optimization steps for each set (OL3, gHBfix-18Ab-pre1, gHBfix-18Ab-pre2, gHBfix-18Ab, and gHBfix-18Ab*) are defined in Table 2 and Figure S3.

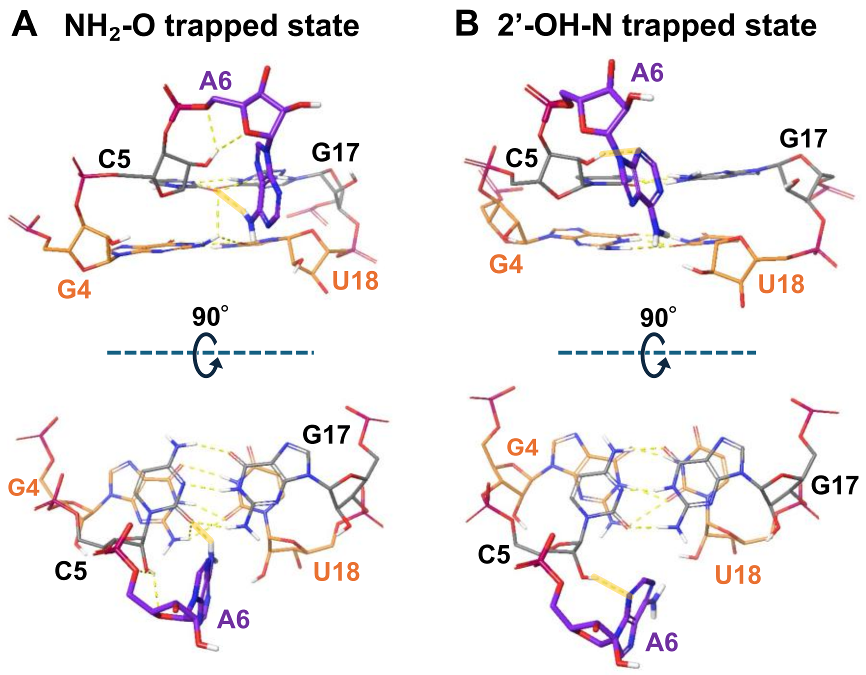


**Figure S5.** Structural artifacts observed with gHBfix-18Ab-pre1 and gHBfix-18Ab-pre2
(A) $NH_2$–O trapped state observed with gHBfix-18Ab-pre1, characterized by excessive stabilization of an A6(N6–H)···C5(O2) hydrogen bond. (B) 2′-OH–N trapped state observed with gHBfix-18Ab-pre2, characterized by excessive stabilization of a U18(O2′–H)···A6(N1) hydrogen bond. These structures associated with PMF minima were selected from the trajectory frames whose θ values were closest to the θ coordinates of the corresponding minima (Figures S3A and S3B). Hydrogen bonds targeted in the subsequent parameter-refinement step are shown as thick yellow lines, while all other hydrogen bonds are depicted as yellow dashed lines.

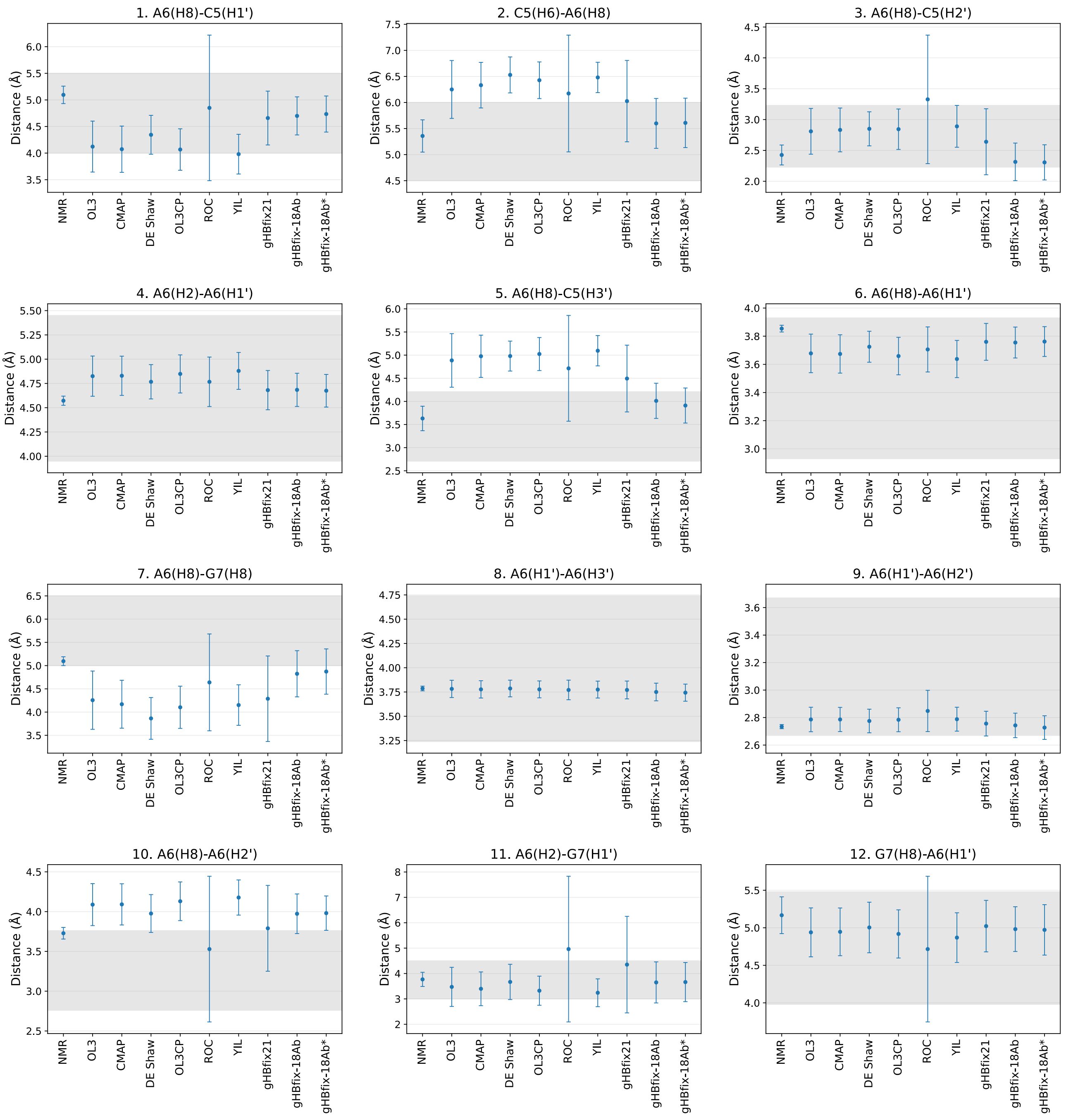


**Figure S6.** Comparison of proton–proton distances around the A6 bulge across the NMR ensemble and AWH simulations

Each panel shows the mean distance and population standard deviation for one of the 12 proton pairs around residue A6. The NMR values were calculated as unweighted averages over the 20 structural models of PDB ID 6VA1. The values for OL3, CMAP, DE Shaw, $OL3_{CP}$, ROC, YIL, gHBfix21, gHBfix-18Ab, and gHBfix-18Ab* are reweighted averages obtained from the corresponding 1000 ns AWH simulations. Error bars indicate population standard deviations, and gray shaded regions represent the experimental NOE distance ranges.

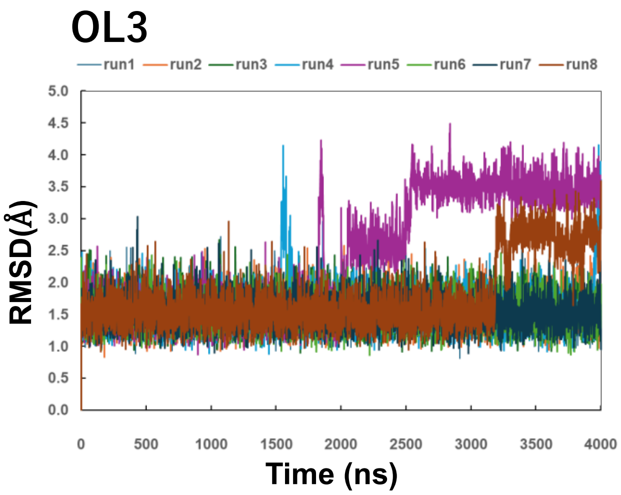


**Figure S7.** RNA all-atom RMSD time series of the cUUCGg tetraloop with OL3

RNA all-atom RMSD values were calculated relative to the initial NMR structure (PDB ID: 2KOC) for eight independent 4 μs conventional MD trajectories initiated with different initial velocities.
